# Endothelial cells promote productive HIV infection of resting CD4+ T cells by an integrin-mediated cell adhesion-dependent mechanism

**DOI:** 10.1101/2020.06.29.177402

**Authors:** Catherine M. Card, Bernard Abrenica, Lyle R. McKinnon, T. Blake Ball, Ruey-Chyi Su

**Author notes:** Corresponding Author (CC).

## Abstract

Resting CD4+ T cells do not support HIV replication *in vitro*, yet are primary targets of early HIV infection events *in vivo*. There is an established role for factors in the tissue microenvironment, including endothelial cells, in enhancing the susceptibility of resting CD4+ T cells to productive infection, yet the mechanisms behind this are not well understood. Endothelial cells facilitate immune cell trafficking throughout the body. Cell adhesion molecules expressed by endothelial cells engage integrins on activated and memory T cells and mediate transmigration into inflamed tissues. These cell trafficking pathways have overlapping roles in facilitating HIV replication but their relevance to endothelial cell-mediated enhancement of HIV susceptibility in resting CD4+ T cells has not previously been examined. We used flow cytometry to characterize the phenotype resting CD4+ T cells that became productively infected when exposed to HIV in the presence of endothelial cells. Infected CD4+ T cells were primarily central memory cells enriched for high expression of the integrins LFA-1 and VLA-4 and had variable expression of α4β7, CCR6 and CD69. Blocking LFA-1 and VLA-4 on resting CD4+ T cells abrogated infection in the co-culture model, indicating that engagement of these integrins is essential for enhancement of resting CD4+ T cell HIV susceptibility by endothelial cells. Cellular activation of CD4+ T cells did not appear to be the primary mechanism enabling HIV replication since only a small proportion of resting CD4+ T cells became activated over the course of the co-culture and fewer than half of infected cells had an activated phenotype. The demonstration that endothelial cells enhance the cellular HIV susceptibility of resting memory CD4+ T cells through cell trafficking pathways engaged during the transmigration of memory T cells into inflamed tissues highlights the physiological relevance of these findings for HIV acquisition and opportunities for intervention.

**Author Summary:** HIV acquisition risk per coital act is relatively low, but this risk is amplified by various behavioural and biological variables. Genital inflammation is a key biological variable associated with increased risk of HIV acquisition, but the mechanisms driving this are incompletely understood. Inflammation is a complex process, with direct effects on HIV target cells as well as the tissue in which those cells reside and encounter virus. The first HIV target cells *in vivo* are resting memory CD4+ T cells, yet these cells are do not support viral replication when purified and exposed to HIV *in vitro*. Rather, signals from tissue microenvironment are required to support viral replication within resting memory CD4+ T cells. Endothelial cells line tissue vasculature and guide immune cell trafficking to inflamed tissues through engagement of integrins by endothelial-expressed cell adhesion molecules. We show here that these same cell-trafficking pathways enable endothelial cells to promote HIV replication within resting memory CD4+ T cells *in vitro*. Blockade of integrins on resting memory CD4+ T cells prevented endothelial enhancement of HIV infection. These findings further our understanding of the determinants of cellular susceptibility to HIV infection and offer a potential mechanism by which inflammation promotes HIV acquisition.

## Introduction

The biological factors that favour HIV acquisition remain incompletely understood. Increased susceptibility to HIV infection has been linked with mucosal inflammation. Specifically, genital inflammation was a strong predictor of HIV acquisition in the CAPRISA 004 trial and inflammation undermined effectiveness of tenofovir gel in prevention of infection [1,2]. Inflammation may promote HIV infection due to impairment of the mucosal barrier [3] and recruitment of activated HIV target cells to mucosal sites [4]. However, inflammation has additional effects on mucosal tissue that may also contribute to increased HIV susceptibility.

HIV preferentially replicates in activated CD4+ T cells, whereas resting CD4+ T cells demonstrate relative resistance to infection, particularly *in vitro* [5-10]. However, non-human primate models have demonstrated that the initial founder population of SIV-infected cells is primarily resting CD4+ memory T cells [11,12]. Furthermore, HIV latency is maintained in the resting memory CD4+ T cell population [11-17], although there is debate about whether this reservoir arises from infection of resting cells or return of infected activated cells to a resting state. These seemingly paradoxical findings may be reconciled by the observation that resting CD4+ T cells become susceptible to HIV infection *in vitro* when cultured in the presence of lymphoid tissue [13], implicating factors in the tissue microenvironment that render resting cells more permissive to HIV infection. Endothelial cells (ECs), which are abundant in lymphoid and mucosal tissues, may partially account for this observation, as they have been shown to enhance productive and latent HIV infection of resting CD4+ T cells [18-22] and are capable of *trans*-infection of CD4+ T cells *in vitro* [23,24]. There is no clear consensus on the mechanisms by which ECs enhance susceptibility of resting CD4+ T cells. In early studies using an allogenic stimulation model, MHC-II on ECs partially contributed to enhancement of infection [18,19], but resting ECs, which lack MHC-II expression, were also shown to enhance infection by cell contact-dependent and - independent mechanisms [20,21]. Among soluble factors tested, IL-6 was implicated as a contributor to enhancement of infection, but other unidentified soluble factors may also play a role under some conditions [21].

ECs are non-hematopoietic cells that line the vessels of the blood and lymphatic systems. Although ECs play multiple roles in shaping immune responses, they are best known for their function as mediators of cell homing to tissues and lymph nodes. This occurs through production of chemokine gradients, which direct cellular migration, and binding of lymphocyte integrins and selectins to their cognate endothelial-expressed cellular adhesion molecule (CAM) ligands. Such EC adhesion molecules include E-selectin, intercellular adhesion molecule 1 (ICAM-1), vascular cell adhesion molecule 1 (VCAM-1) and mucosal vascular addressin cell adhesion molecule 1 (MAdCAM-1). Under inflammatory conditions, cytokines such as TNFα and IL-1β produced by immune and stromal cells in tissue lead to changes in the vascular endothelium, including upregulation of VCAM-1 and ICAM-1, thereby promoting recruitment of T cells bearing their cognate ligands, LFA-1 (αLβ2) and VLA-4 (α4β1), respectively.

Cell homing pathways have previously been implicated in HIV susceptibility and pathogenesis. Expression of LFA-1 integrin on CD4+ T cells promotes HIV entry through interactions between LFA-1 and its ligand, ICAM-1, incorporated into the virion envelope [25-27]. ICAM-1 and LFA-1 have also been implicated in cell-cell spread of HIV, in which adhesion between donor and target cells is a critical step in establishing the virological synapse [28,29]. Consistent with the role of this pathway in promoting HIV infection, inhibition of interactions between LFA-1-and ICAM-1 reduce cell-cell spread of HIV and infection by ICAM-bearing viruses *in vitro* [25,30]. HIV preferentially targets CD4+ T cells expressing VLA-4 and the gut-homing integrin α4β7 [31-37]. α_4_β_7_ has been shown to provide co-stimulation to TCR-activated CD4+ T cells when engaged by MAdCAM-1, thereby promoting productive HIV replication *in vitro* [38]. Blocking of α_4_β_7_ in monkey models showed promise for prevention of SIV infection and for viral control in SIV-infected monkeys [39,40]. However, the latter results were not substantiated in recent studies of non-human primates [41-43] or HIV-infected patients [44].

Given the relevance of cell homing pathways to HIV pathogenesis and the role of ECs in regulating cell trafficking, we hypothesized that molecules that function in homing pathways may also mediate the enhancement of HIV infection observed when resting CD4+ T cells are cultured with ECs. To address this gap in knowledge, we cultured resting CD4+ T cells in the presence of ECs, and subsequently infected them with HIV *in vitro*. We show here that resting CD4+ T cells expressing high levels of integrins LFA-1 and VLA-4 are preferentially infected. Targeted blocking of these integrins prevented adhesion of CD4+ T cells to ECs and diminished the EC-mediated promotion of HIV infection.

## Results

### ECs enhance infection of resting CD4+ T cells

We first sought to confirm previous reports indicating that resting CD4+ T cells (rCD4) are susceptible to productive HIV infection when co-cultured with ECs [18-22]. Fresh rCD4 cells were isolated from peripheral blood mononuclear cells (PBMC) from healthy donors. Magnetic negative isolation resulted in a rCD4 population that was CD3+ CD4+ HLA DR-CD25-/dim CD69- (median purity 97.2% of total CD45+ cells, Fig 1). rCD4 cells were further immunophenotyped to determine baseline expression of markers of memory (CD45RA, CCR7), activation (CD69, Ki67), integrins VLA-4 (β1), LFA-1 (β2) and α4β7 (β7) and CCR6 on rCD4 and compare to bulk CD4+ T cells in PBMC (Table 1, S1 Fig).

**Table 1.** Comparison of phenotypes between *ex vivo* PBMC and isolated resting CD4+ T cells.

| Parameter <sup>a</sup> | PBMC | rCD4 | p-value <sup>b</sup> |
| --- | --- | --- | --- |
| CD4+ T cell populations |  |  |  |
| CD69 (%CD4+) | 0.37 (0.17-0.80) | 0.06 (0.03-0.42) | **0.020** |
| Ki67 (%CD4+) | 0.44 (0.18-0.93) | 0.086 (0.05-0.45) | **0.002** |
| CD45RA (%CD4+) | 53.9 (45.0-59.5) | 65.5 (59.1-75.7) | **0.004** |
| Naïve CD4+ T cell populations |  |  |  |
| β1<sup>lo</sup> (%CD45RA+) | 99.4 (99.3-99.5) | 99.6 (99.4-99.8) | 0.094 |
| β1 MFI | 794.0 (725.5-742.5) | 853.0 (742.5-895.5) | 0.844 |
| β2<sup>+</sup> (%CD45RA+) | 99.7 (99.4-99.9) | 99.9 (99.8-100.0) | 0.125 |
| β2 MFI | 4681 (1412-6807) | 5552 (1778-5858) | 0.098 |
| β7<sup>int</sup> (%CD45RA+) | 50.2 (38.7-61.4) | 54.3 (43.3-61.7) | **0.043** |
| β7 MFI | 933.0 (437.0-1058) | 944.0 (452.5-1058) | 0.570 |
| CCR7 (%CD45RA+) | 97.7 (94.7-99.1) | 99.3 (98.8-99.9) | **0.004** |
| Memory CD4+ T cell populations |  |  |  |
| β1<sup>hi</sup> (%CD45RA-) | 38.8 (35.1-48.1) | 52.9 (48.6-58.3) | **0.027** |
| β1 MFI | 2068 (1874-2396) | 2021 (1912-2596) | 0.250 |
| β1<sup>lo</sup> (%CD45RA-) | 44.4 (38.3-48.7) | 38.7 (36.4-48.3) | 0.551 |
| β1 MFI | 742.0 (652.5-812) | 720.0 (628.5-853.0) | 0.320 |
| β2<sup>hi</sup> (%CD45RA-) | 99.8 (99.3-100.0) | 99.8 (99.8-100.0) | 0.094 |
| β2 MFI | 7749 (2526-9559) | 8053 (2981-9328) | 0.301 |
| β7<sup>hi</sup> (%CD45RA-) | 18.7 (14.9-27.1) | 25.6 (20.9-31.6) | **0.004** |
| β7<sup>hi</sup> MFI | 3450 (1762-3771) | 3388 (1971-3733) | 0.129 |
| CCR6 (%CD45RA-) | 30.5 (15.3-38.4) | 40.2 (31.2-49.2) | **0.039** |
| CCR6 MFI | 1884 (1442-2640) | 2433 (2230-2824) | **0.012** |
| CCR7 (%CD45RA-) | 74.3 (69.7-76.6) | 87.7 (84.8-91.7) | **0.004** |

<sup>a</sup> expression values listed as median of % positive (+) cells in indicated parent population or median fluorescence intensity (MFI), with interquartile range shown in parentheses
<sup>b</sup> p-values calculated using non-parametric Wilcoxon matched pairs test. Parameters considered statistically significant (p<0.05) are highlighted in bold text

**Figure 1.**
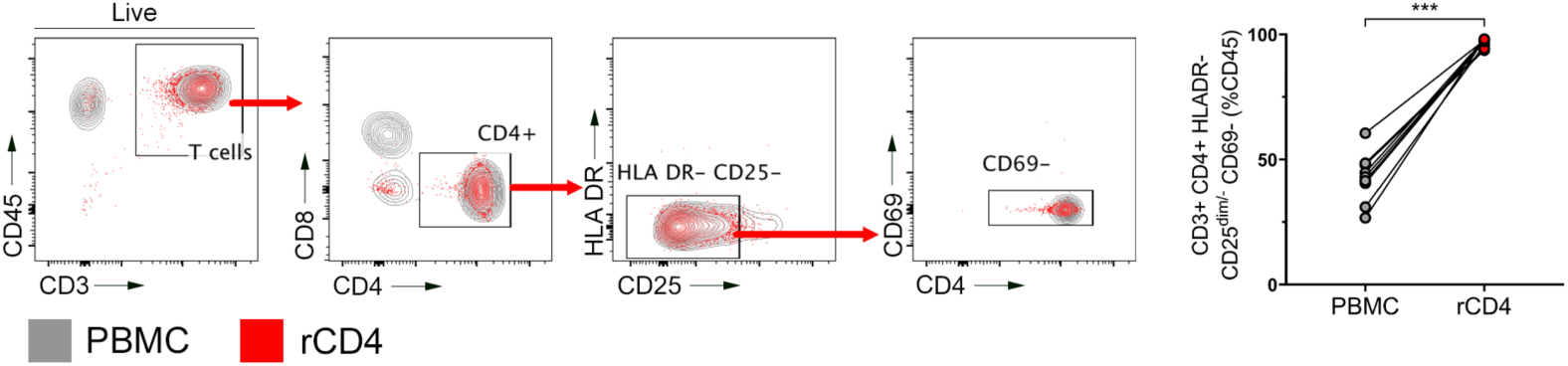
Phenotype of resting CD4+ T cells (rCD4) following isolation. Representative flow cytometry data showing gating of CD4+ HLA DR-CD25dim/-CD69-T cells in PBMC (grey) or isolated rCD4 cells (red) (left panel). Cells were gated on live lymphocytes then were analyzed for the indicated markers. Quantification of the purity of resting CD4+ T cells in the indicated sample types (right panel). Statistical comparison was performed by Wilcoxon matched-pairs signed-rank test (*** p<0.001).

Compared to bulk CD4+ T cells, a greater proportion of isolated rCD4 cells were naïve, as indicated by CD45RA expression (median 53.9% vs. 65.5%, respectively; p=0.004). This was expected, as naïve cells, by definition, have not previously been activated by their cognate antigen are therefore expected to be quiescent. Among CD45RA+ naïve populations, greater proportions of rCD4 cells expressed integrin β7 (54.3% vs 50.2%, p=0.043) and CCR7 (99.3% vs 97.7%, p=0.004) compared to bulk CD4+ T cells in PBMC. Among CD45RA-memory cells, greater proportions of rCD4 cells were CCR7+ (87.7% vs 74.3%, p=0.004), indicating a central memory phenotype. rCD4 cells were also enriched for integrin β1^hi^ (42.9% vs. 38.8%, p=0.027), integrin β7^hi^ (25.6% vs 18.7%, p=0.004) and CCR6+ (40.2% vs. 30.5%, p=0.012) populations compared to bulk CD4+ T cells in PBMC. CCR6 receptor density was higher on CCR6+ rCD4 than CCR6+ bulk CD4+ T cells (MFI 2433 vs. 1884, p=0.012). Although expression of activation markers was low in *ex vivo* PBMC, fewer isolated rCD4 expressed the early activation marker CD69 (0.06% vs. 0.37%, p=0.02) or the cell cycle marker Ki67 (0.086 vs. 0.44%, p=0.002) compared to bulk CD4+ T cells, consistent with depletion of activated cells from PBMC (Table 1, S1 Fig).

Isolated rCD4 cells were co-cultured with resting or activated (TNFα-treated) ECs for 24 hours prior to exposure of co-cultures to HIV_IIIB_. Three days post-infection, culture supernatants and cells were collected to assess frequencies of HIV-infected cells and cell phenotypes (Fig 2A). As expected, productively-infected cells were not detectable when rCD4 cultured alone were exposed to HIV (Fig 2B). However, consistent with previous work [18-22], co-culture with EC or TNFα-treated EC resulted in a significant increase in the proportion of productively-infected cells (p=0.0002), indicated by intracellular staining of HIV-p24 protein (range 2.86-30.0% and 2.63-26.4% for HUVEC and HUVEC-TNFα, respectively; Fig 2B). Consistent with these observations, significantly more HIV-p24 protein was detectable in culture supernatants from rCD4 cells co-cultured with TNFα-treated EC compared to those cultured alone (Kruskal-Wallis p = 0.004, rCD4 alone vs. HUVEC-TNFα Dunn’s post-test p=0.0003).

**Figure 2.**
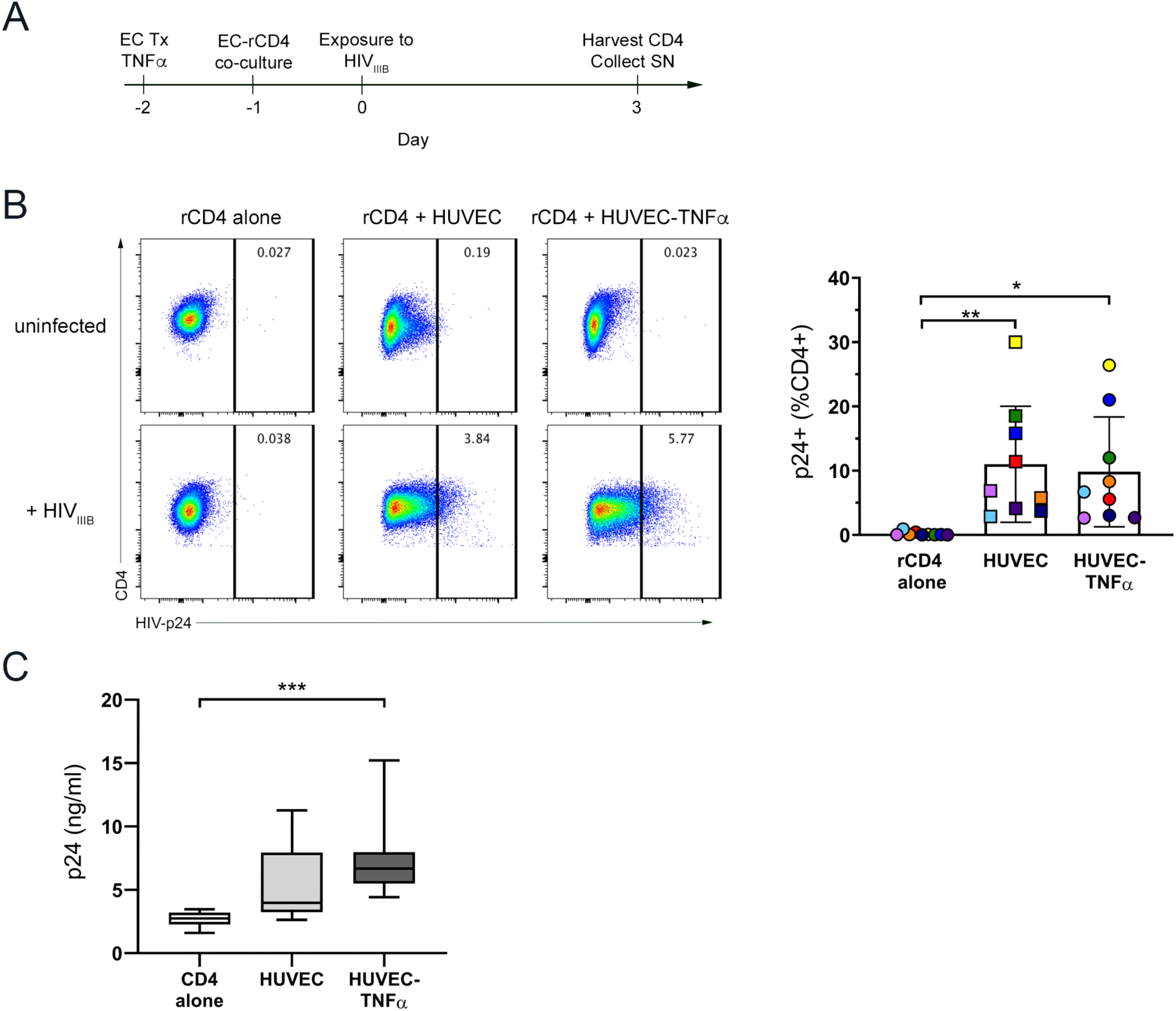
ECs promote HIV infection of rCD4. (A) Schematic timeline of EC-rCD4 co-culture infection assay. (B) Representative flow cytometry data demonstrating intracellular detection of HIV-p24 protein in rCD4 cells cultured alone or in the presence of untreated or TNFα-treated HUVECs. Cells were gated on live, CD45+ CD4+ lymphocytes then were analyzed for HIV-p24. Graph shows quantification of flow cytometry data. Data points indicate different CD4+ T cell donors and each donor is indicated by a different colour. Co-culture of rCD4 with HUVEC or HUVEC-TNFα resulted in a significant increase in the proportion of rCD4 cells that became productively infected with HIV (Friedman p=0.0002). (C) Quantification of HIV-p24 protein in cell culture supernatants by HIV-p24 ELISA. Co-culture of rCD4 with HUVEC-TNFα resulted in a significant increase in levels of supernatant HIV-p24 (Kruskal-Wallis p=0.0004). Statistics assessed by Friedman or Kruskal-Wallis test and Dunn post-test (* p<0.05, ** p < 0.01, *** p<0.001 in post-test comparisons).

### High expression of integrins β1 (VLA-4) and β2 (LFA-1) marks HIV-susceptible cells in EC co-cultures

We further examined markers of memory (CD45RA, CCR7), activation (CD69, Ki67), integrins VLA-4 (β1), LFA-1 (β2) and α4β7 (β7) and CCR6 on infected and uninfected rCD4 cells from co-cultures with TNFα-treated ECs (Fig 3A). The majority of infected (HIV-p24+) cells had a central memory phenotype (CD45RA-CCR7+; 52.5%). This was in contrast to uninfected (HIV-p24-) cells, which were CD45RA+ CCR7+ (87.2%), consistent with a naïve phenotype (HIV-p24+ vs HIV-p24-p=0.004; Fig 3B and S1 Table). Infected cells were enriched for high expression of integrins VLA-4 (as indicated by β1 expression), LFA-1 (β2) and β7 as well as CCR6 and the activation marker CD69 (all p=0.004; Fig 3B and S1 Table). Although cells expressing the proliferation marker Ki67 were also enriched in the infected cell population (p=0.004; Fig 3B and S1 Table), they were relatively rare (range 1.71-3.11% of HIV-p24+ cells).

**Figure 3.**
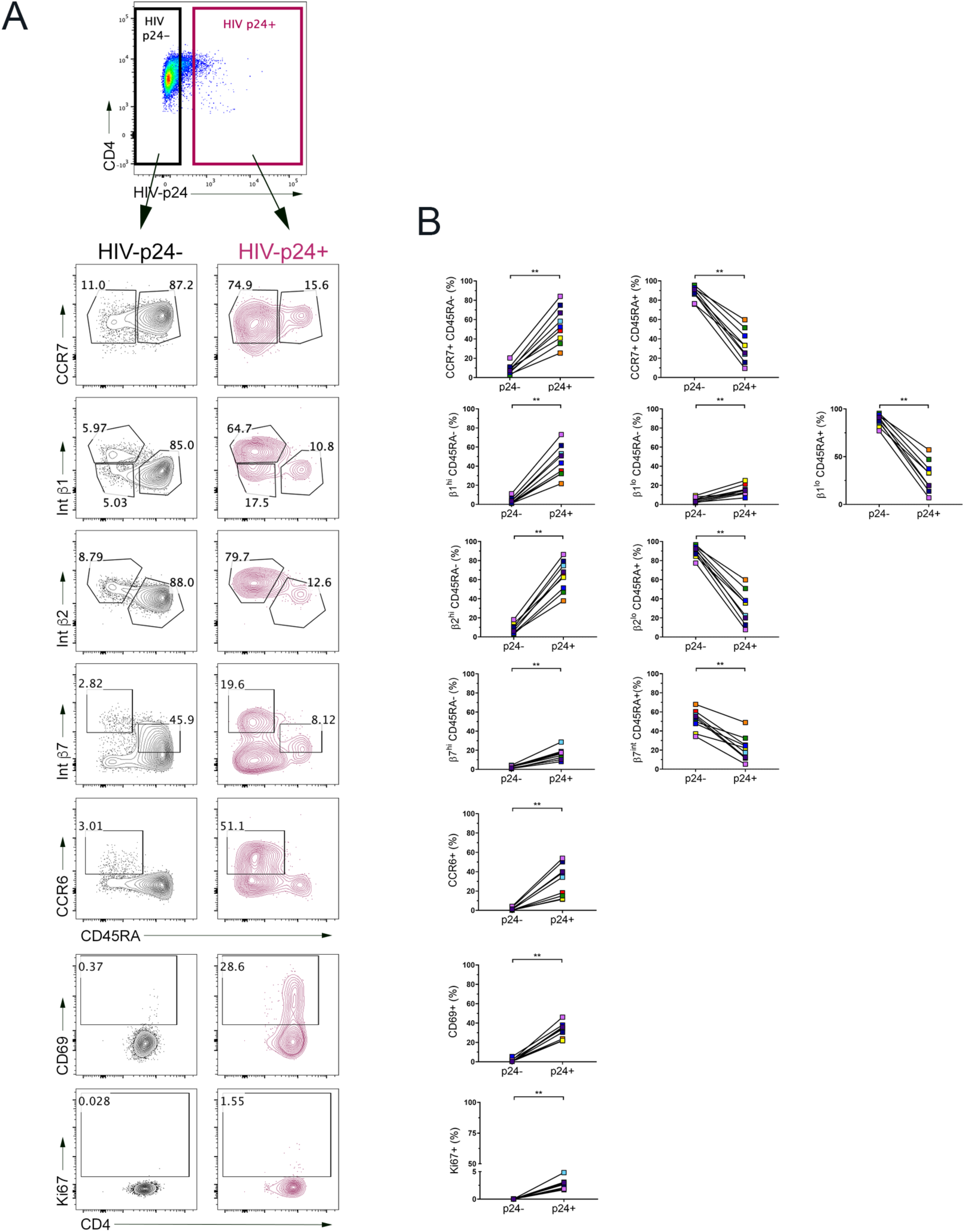
Phenotypic comparison of HIV-infected and uninfected rCD4. (A) Representative flow cytometry showing characterization of HIV-uninfected (HIV-p24-, grey) and HIV-infected (HIV-p24+, purple) rCD4 cells from co-cultures with TNFα-treated ECs assessed for markers of activation (CD69, Ki67), memory (CD45RA, CCR7), integrins VLA-4 (β1), LFA-1 (β2) and α4β7 (β7) and CCR6. (B) Quantification and comparison of flow cytometry plots in (A). Statistical comparisons were performed by Wilcoxon matched-pairs signed-rank test (** p < 0.01).

We next evaluated whether these were exclusive markers of different HIV-susceptible subsets cells or whether the same subset of HIV-susceptible cells co-expressed multiple enriched markers. To achieve this, a Boolean gating strategy was applied to the populations with the highest expression of each enriched marker (Fig 4A) in both infected (HIV-p24+) and uninfected (HIV-p24-) subsets from rCD4 cells infected in the presence of TNFα-treated ECs. As Ki67 expression was low on both infected and uninfected cell subsets, it was excluded from co-expression analysis. The infected cell population was heterogenous and comprised of numerous subsets defined by combinations of the specified markers (Fig 4A). There was also a subset of cells that did not express any of the specified markers, which was highly enriched among the uninfected cells (median 28.6% of HIV-p24+ vs 92.47% of HIV-p24-cells; p=0.004). Despite this heterogeneity, three main phenotypic groupings were evident based on expression of β_1_ and β_2_ integrins. Cell subsets that were β1^hi^ β2^hi^ and β1^lo/-^ β2^hi^ were enriched among infected compared to uninfected cells, and had variable expression of β7, CCR6 and CD69 (Fig 4A). A third phenotype that was overrepresented among infected cells was marked solely by CD69 expression in the absence of high levels of β1 and β2 integrins (Fig 4A). These subsets were clearly represented when we repeated the co-expression analysis to focus specifically on β1, β2 and CD69 (Fig 4B). Specifically, cells that co-expressed high levels of β1 and β2 integrins, either in the presence or absence of CD69, accounted for approximately half of infected cells (median β1^hi^ β2^hi^ CD69+ = 24.4%, median β1^hi^ β2^hi^ CD69-= 22.8% of HIV-p24+) but fewer than 2% of uninfected cells (median β1^hi^ β2^hi^ CD69+ = 0.13%, median β1^hi^ β2^hi^ CD69-= 1.65% of HIV-p24-; HIV-p24+ vs HIV-p24-p=0.0003, p=0.0005 for CD69+ and CD69-subsets, respectively; Fig 4B). The infected cell population was further enriched for β1^lo/-^ β2^hi^ CD69+ cells and other minor subsets, including cells that expressed CD69 in the absence of high levels of β1 and β2 integrins (Fig 4B). Collectively, these data implicate β1^hi^ β2^hi^ and β1^lo/-^ β2^hi^ cells as the primary targets for HIV infection in the presence of endothelial cells, with CD69+ cells lacking high expression of integrins as an additional HIV target.

**Figure 4.**
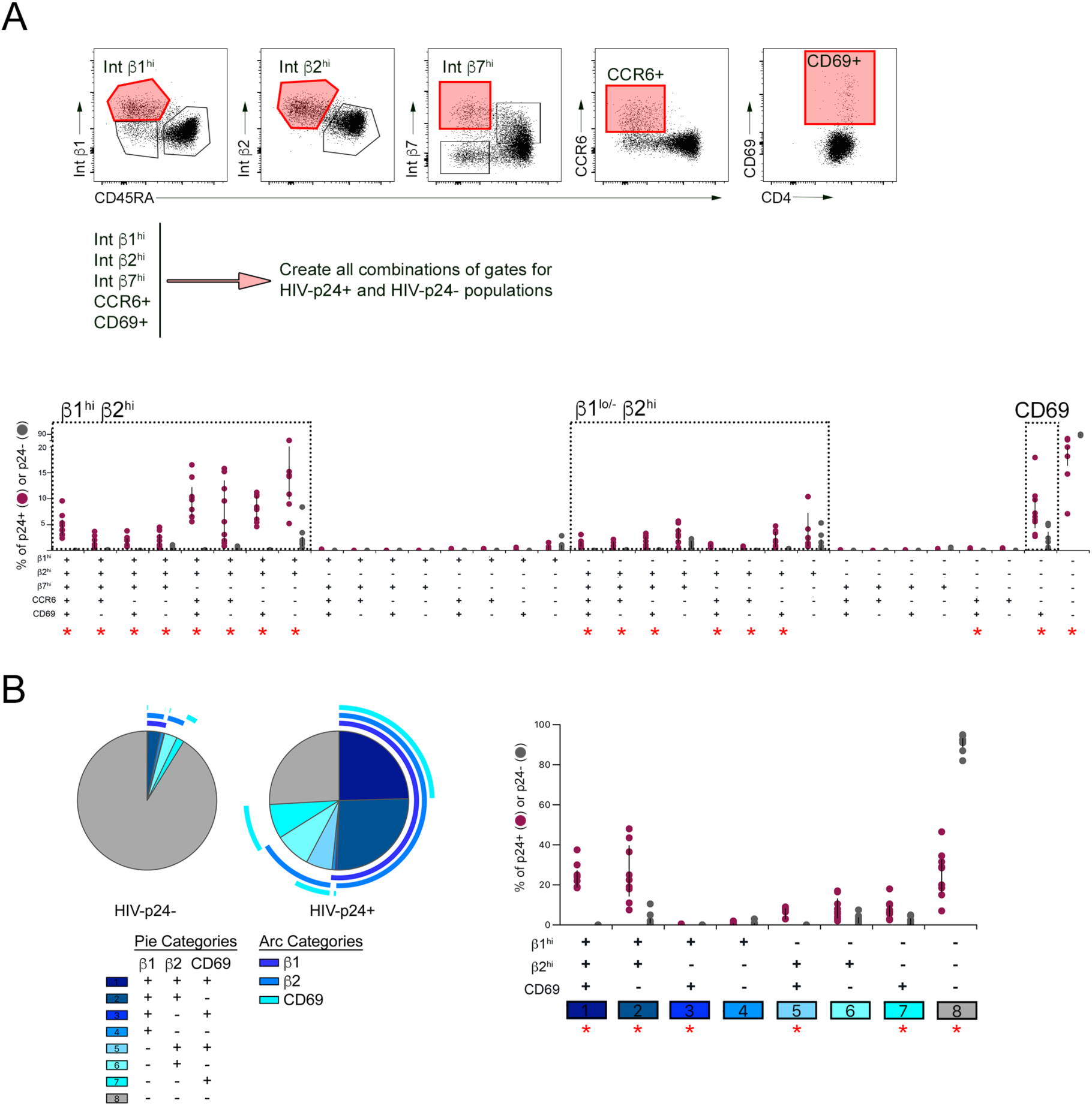
Co-expression analysis of phenotypic markers in HIV-infected and uninfected rCD4. (A) A Boolean gating strategy was applied to the populations with the highest expression of integrins β1, β2 and β7, CCR6 and CD69, indicated by the red gates in flow cytometry plots (top panel). Graph shows comparison of HIV-infected (HIV-p24+, purple circles) and uninfected (HIV-p24-, grey circles) cells for each phenotypic combination (bottom panel). (B) Co-expression analysis of integrins β1 and β2 and CD69 among HIV-infected and uninfected cells. Pie graphs indicate relative contribution of each combination to uninfected (HIV-p24-) or infected (HIV-p24+) population. Each marker is indicated by an arc surrounding the graph. Overlapping arcs indicate co-expression. Graph shows comparison of HIV-infected (HIV-p24+, purple circles) and uninfected (HIV-p24-, grey circles) cells for each phenotypic combination. Co-expression of phenotypic markers on uninfected (HIV-p24-) and HIV-infected (HIV-p24+) CD4+ T cells was performed using SPICE software. Statistical comparisons were performed by Wilcoxon matched-pairs signed-rank test. Red asterisks indicate significant comparisons after Bonferroni correction (p < 0.002 for A, p< 0.006 for B).

### Endothelial cells express ligands for integrins LFA-1 (αLβ2) and VLA-4 (α4β1)

Integrins LFA-1 (αLβ2) and VLA-4 (α4β1) and are known to engage endothelial-expressed cell adhesion molecules ICAM-1 and VCAM-1, respectively, during cellular transmigration into inflamed tissues. To confirm that these ligands were expressed on the HUVECs used as ECs in this co-culture model, HUVECs were analyzed by flow cytometry for surface ICAM-1 and VCAM-1, as well as MAdCAM-1, the ligand for integrin α4β7, under steady state and inflammatory (TNFα-treated) conditions. HLA DR (MHC class II) was also measured to address whether HUVECs might be able to activate rCD4 via allo-stimulation, as the primary HUVECs and rCD4 cells were isolated from different donors. IFNγ treatment of ECs was included as a positive control for upregulation of HLA DR expression. ICAM-1 was moderately expressed under steady state conditions and was upregulated following TNFα stimulation (Fig 5A). VCAM-1 was similarly upregulated following treatment with TNFα and the VCAM-1+ population co-expressed high levels of ICAM-1 (Fig 5B). As expected, HUVECs were negative for both MAdCAM-1 and HLA DR at both steady state and following treatment with TNFα, whereas HLA DR was upregulated by IFNγ stimulation (Fig 5A).

**Figure 5.**
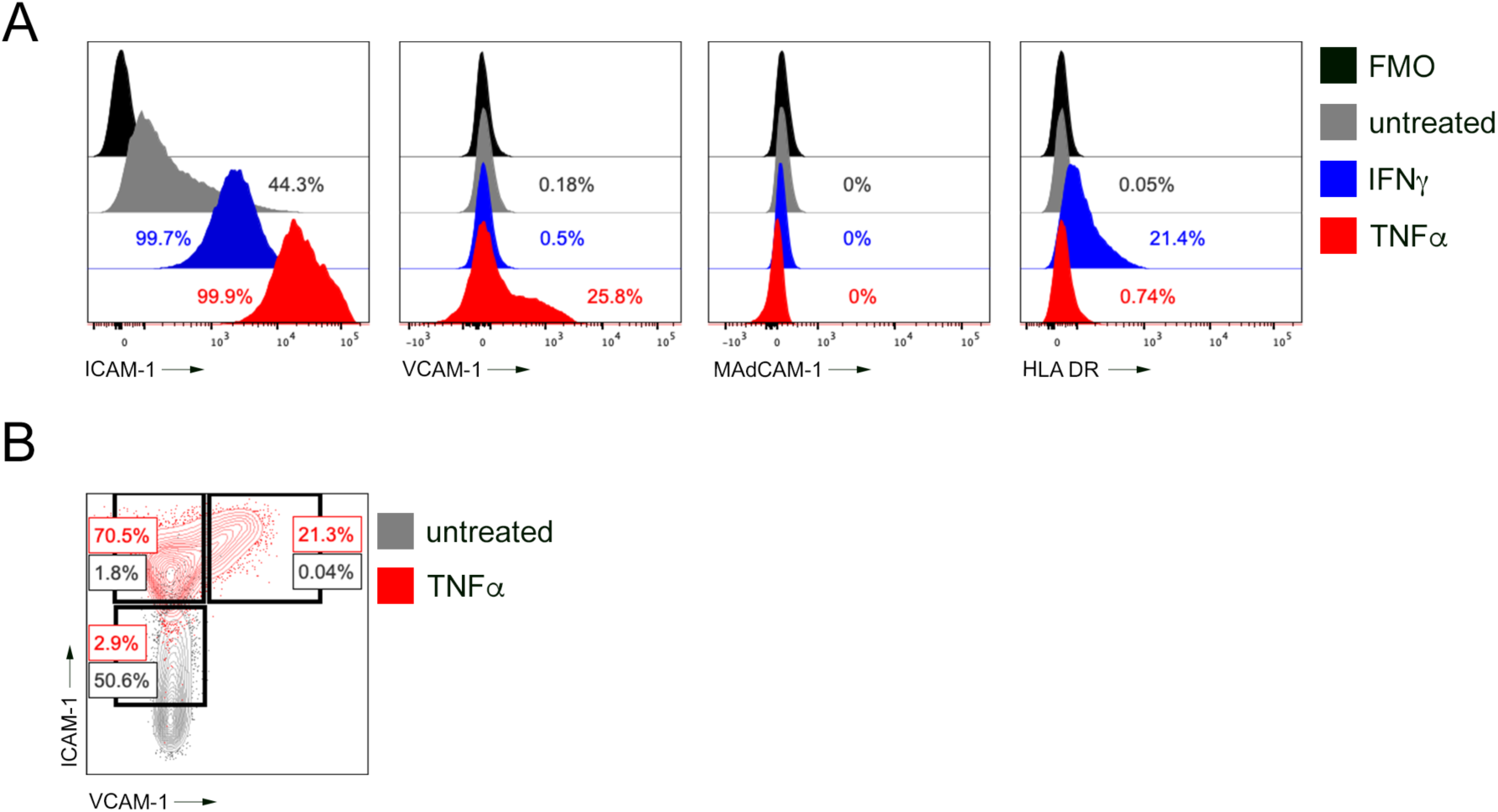
Cell adhesion molecules are expressed by ECs and upregulated under inflammatory conditions. (A) Representative flow cytometry data demonstrating expression of ICAM-1, VCAM-1, MAdCAM-1 and HLA DR on untreated (grey histograms) TNFα-treated (red histograms) or IFNγ-treated (blue histograms) HUVECs. Cells were first gated on live, CD45-CD31+ cells then were analyzed for the different phenotypic markers using fluorescence minus one (FMO) controls (black histograms). (B) Representative flow cytometry analysis of ICAM-1 and VCAM-1 co-expression on untreated (grey) or TNFα-treated (red) HUVECs.

### EC-mediated enhancement of HIV infection of rCD4 cells is dependent on integrins

To determine whether engagement of integrins by ECs was necessary for enhancement of rCD4 infection, we used integrin-specific blocking antibodies to interfere with this interaction. We elected to use integrin α-chain-specific antibodies with the aim of preserving the ability to detect the integrin β chains by flow cytometry following blocking. Antibody clones TS1/22 (anti-LFA1 αL chain) and 2B4 (anti-VLA-4 α4 chain) were selected based on previous reports of blocking capability [45,46]. Consistent with this, incubation of rCD4s with anti-LFA-1 (TS1/22) and anti-VLA-4 (2B4) inhibited adhesion to both TNFα-treated HUVECs and recombinant ICAM-1 or VCAM-1-coated plates in a fluorescence-based static adhesion assay, compared to isotype control (mouse IgG1κ, clone 11711, S2 Fig). Unfortunately, blocking of α4 did interfere with staining of integrins VLA-4 (β1 chain) and α4β7 (β7 chain) but blocking of αL did not have any effect on the detection of LFA-1 (β2 chain) (S3 Fig). This interference of β-chain detection precluded further characterization of integrin expression on cells after blocking treatments.

Blockade of LFA-1 and VLA-4 in the co-culture infection model (Fig 6A) inhibited HIV infection of rCD4s co-cultured with TNFα-treated ECs, whereas isotype control antibody had no effect (p=0.0002, Fig 6B). Consistent with these flow cytometry results, detection of HIV-p24 protein was significantly reduced in supernatants collected from co-cultures with LFA-1 and VLA-4 blockade (p=0.0007, Fig 6C).

**Figure 6.**
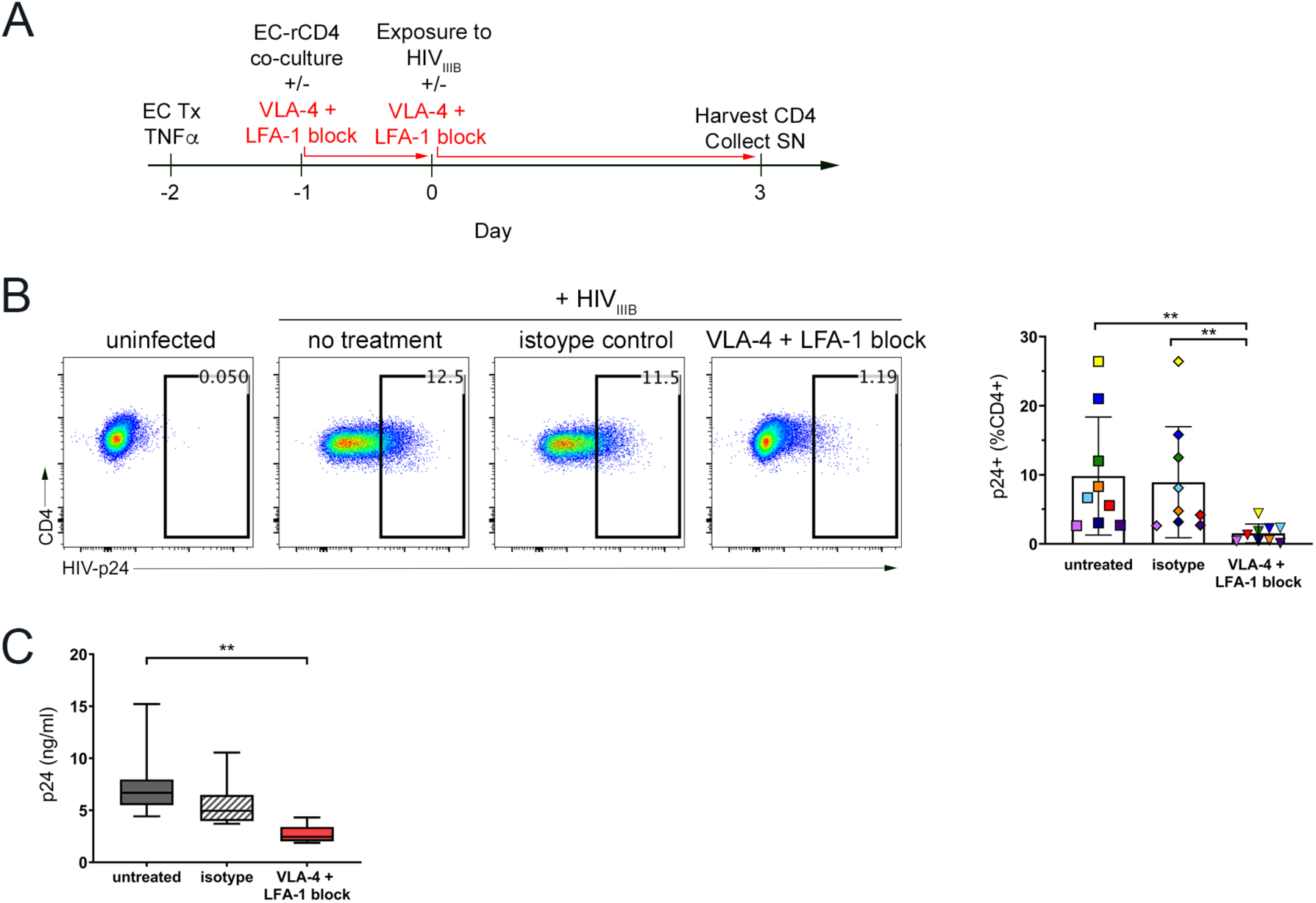
Integrin blockade prevents EC-mediated enhancement of rCD4 infection by HIV. (A) Schematic timeline of EC-rCD4 co-culture infection assay. (B) Representative flow cytometry data showing intracellular detection of HIV-p24 protein in rCD4 cells co-cultured with ECs in the absence of treatment or after treatment with anti-LFA-1 (αL chain, clone TS1/22) and anti-VLA-4 (α4 chain, clone 2B4) or isotype control antibody (clone 11711). Cells were gated on live, CD45+ CD4+ lymphocytes then were analyzed for HIV-p24. Graph shows quantification of flow cytometry data. Integrin blockade resulted in a significant decrease in the proportion of rCD4 cells that became productively infected with HIV (Friedman p=0.0002). (C) Quantification of HIV-p24 protein in cell culture supernatants by HIV-p24 ELISA. Integrin blockade resulted in a significant decrease in levels of supernatant HIV-p24 (Friedman p=0.0007). Statistics assessed by Friedman test with Dunn post-test (** p < 0.01 in post-test comparisons).

### Endothelial engagement of rCD4 cells may promote infection by inducing low level cellular activation

In contrast to resting cells, activated CD4+ T cells readily support productive HIV replication *in vitro*. To determine if ECs promoted productive HIV infection in rCD4s by inducing cellular activation, rCD4s were co-cultured with ECs in the absence of HIV for four days (equivalent to total co-culture time in infection assays) then cellular activation was determined by CD69 and Ki67 expression. Significantly higher proportions of rCD4s co-cultured with ECs expressed CD69 and Ki67 than those cultured alone (Fig 7A). In contrast, rCD4 cells cultured alone remained resting, with similar proportions of CD69- and Ki67-expressing cells as freshly-isolated rCD4 (S2 Table).

**Figure 7.**
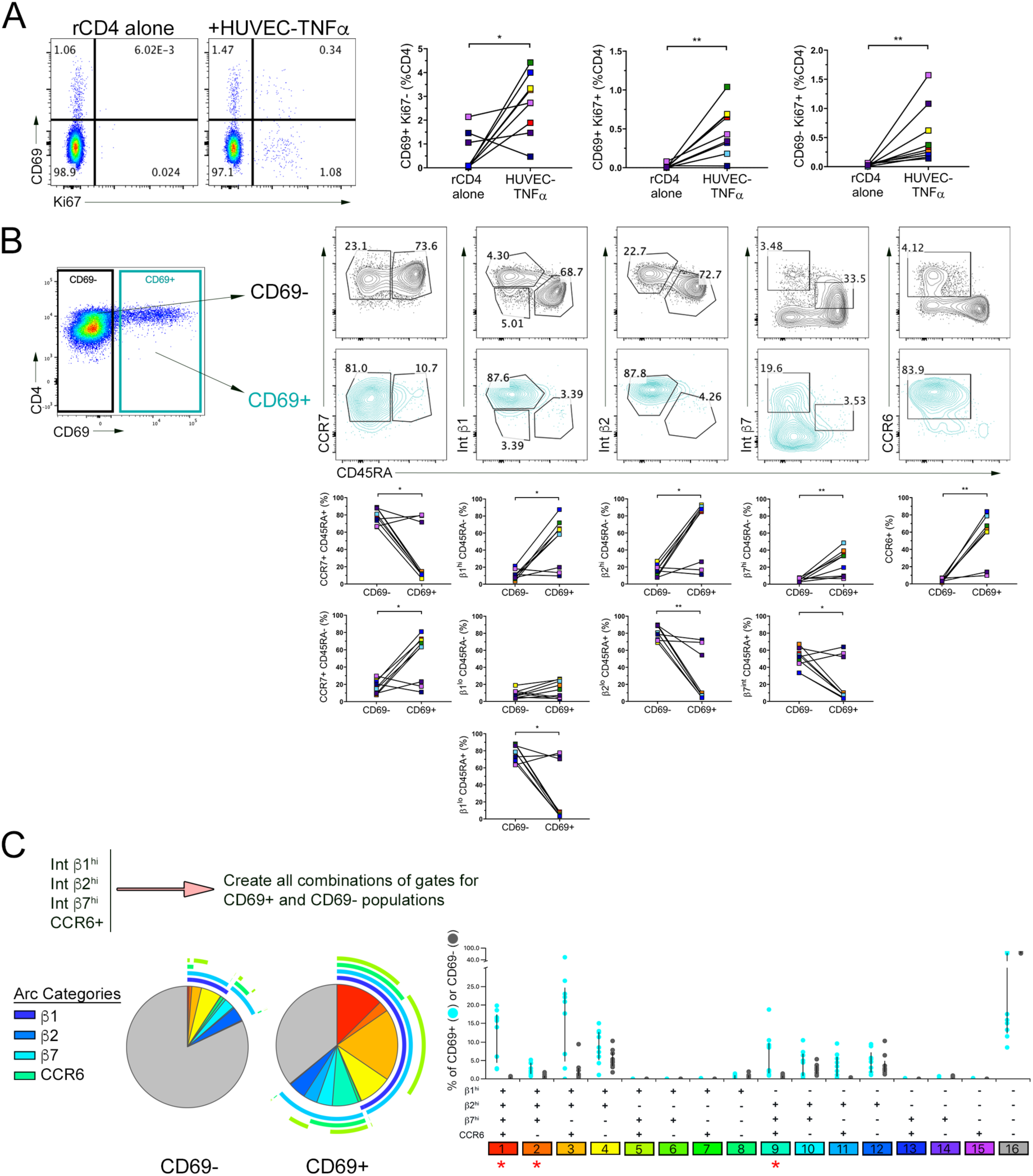
ECs induce cellular activation of rCD4. (A) Analysis of CD69 and Ki67 on rCD4 following co-culture with ECs. Statistical comparisons were performed by Wilcoxon matched-pairs signed-rank test (* <0.05, ** p < 0.01). (B) Resting (CD69-, black contour plots) and activated (CD69+, blue contour plots) CD4+ T cells from EC co-cultures were assessed for memory markers (CD45RA, CCR7), integrins VLA-4 (β1), LFA-1 (β2) and α4β7 (β7) and CCR6. Graphs show quantification and comparison of flow cytometry plots. Statistical comparisons were performed by Wilcoxon matched-pairs signed-rank test (* <0.05, ** p < 0.01). (C) Co-expression of β1^hi^, β2^hi^, β7^hi^ and CCR6+ phenotypes within resting (CD69-) and activated (CD69+) CD4+ T cell populations. Pie graphs indicate relative contribution of each combination to resting (CD69-) or activated (CD69+) population. Each marker is indicated by an arc surrounding the graph. Overlapping arcs indicate co-expression. Graph shows comparison of activated (CD69+, blue circles) and resting (CD69-, grey circles) cells for each phenotypic combination. Co-expression analysis was performed using SPICE software. Statistical comparisons were performed by Wilcoxon matched-pairs signed-rank test. Red asterisks indicate significant comparisons after Bonferroni correction (p < 0.004).

CD69-expressing cells were further characterized for expression of CD45RA, CCR7, VLA-4 (β_1_), LFA-1 (β_2_), integrin β7 and CCR6. Consistent with the phenotype observed for HIV-p24+ cells, CD69+ cells were primarily central memory cells (CD45RA-CCR7+), enriched for high expression of integrins VLA-4 (β1), LFA-1 (β2) and variable expression of β7. CCR6 was also frequently expressed among CD69+ cells (Fig 7B). Furthermore, there was a high level of co-expression between these markers, with significant enrichment of cell subsets co-expressing high levels of β1, β2 and β7 integrins in the presence or absence of CCR6 within the CD69+ vs CD69-cell population (Fig 7C). Collectively, these data suggest that TNFα-treated ECs induce some activation of memory rCD4s that may help support HIV replication. However, only a small proportion of rCD4s expressed CD69 after three days of co-culture with ECs (median 3.3%). Indeed, in matched samples, there were significantly more rCD4 cells that became infected than those that became activated when co-cultured with ECs (p=0.008; S4 Fig). Furthermore, as shown in Fig 3A and S1 Table, fewer than half of HIV-p24+ cells expressed CD69 (median 33.8%), suggesting that cellular activation is not the main mechanism driving enhanced HIV replication in the presence of ECs.

## Discussion

Resting CD4 cells are known to be early HIV targets *in vivo*, in contrast to their relative resistance to infection *in vitro*. There is an established role for the tissue microenvironment, and ECs in particular, in enhancing HIV susceptibility of rCD4 cells. However, the mechanisms by which this occurs are not well understood. In this study we have demonstrated that in the presence of ECs, HIV infection of rCD4 cells occurs primarily in a subset of memory cells expressing high levels of integrins LFA-1 and VLA-4. ICAM-1 and VCAM-1, the ligands for LFA-1 and VLA-4 respectively, were expressed by TNFα-treated ECs. Blocking of these integrins prevented HIV infection in our co-culture model, indicating that EC-mediated enhancement of HIV infection of rCD4 cells occurs in an integrin-dependent manner.

Adhesion molecules have previously been implicated in promoting HIV infection of CD4+ T cells in multiple ways. HIV is well-known to incorporate host proteins into the vial envelope when budding from a cell. In particular, ICAM-1 [47,48] and α4β7 [49] have been shown to be present on various strains of HIV. HIV virions bearing host-derived adhesion molecules can directly bind their cognate ligands on target cells to promote viral entry and facilitate early events in HIV replication [27,50]. The virus used in our study was propagated in physiologically-relevant human PBMCs and can be expected to bear host proteins, including integrins and CAMs. It is possible that virion-embedded host ICAM-1 could engage LFA-1 on target resting memory CD4+ T cells to facilitate infection. However, infection was not observed in the same resting memory CD4+ T cells in the absence of ECs, demonstrating that infection of resting cells occurs in an EC-dependent manner in our system. Furthermore, preferential HIV entry into α4β7 and VLA-4-expressing cervical CD4+ T cells was previously shown to occur independently of direct viral binding to integrins [31].

ICAM and LFA-1 are also critically important in cell-to-cell spread of HIV. This may occur between an uninfected target CD4+ T cell and either a homotypic infected CD4+ T cell or an alternate cell type that is infected or has captured HIV virions. In the latter case, HIV is captured by cells, such as dendritic cells, via various potential receptors without necessarily becoming productively infected, then the virus is presented *in trans* to a susceptible target CD4+ T cell [28,29]. *Trans*-infection is not limited to DCs, and has been described for ECs [23,24] and other non-hematopoietic cell types including epithelial cells [51,52], fibroblasts [51,53] and lymph node fibroblastic reticular cells [54]. In the present study, it is possible that ECs promote infection of resting memory CD4+ T cells through *trans*-infection by capturing integrin-decorated HIV virions via ICAM or VCAM proteins and transferring them to target cells. LFA-1 / ICAM-1 interactions may further contribute to this process through stabilization of cell-cell interactions [29]. Further work is needed to elucidate the relative contribution of *trans*-infection to enhancement of resting CD4+ T cell infection by ECs in this model.

Homotypic transfer of virus occurs between an infected CD4+ T cell and an uninfected bystander CD4+ T cell. In this case, viral and host proteins cluster at the surface of the infected CD4+ T cell. Interaction with the uninfected target CD4+ T cell is facilitated by ICAM-1 / LFA-1 interactions [55], and viral budding occurs at the site of cell-cell contact by fusion with the target cell membrane [28,29]. Our data do not discount the possibility that the reduction in productive infection observed upon integrin blockade was due, in part, to LFA-1-mediated homotypic cell-to-cell spread following initial infection. After initial infection events, HIV amplification in culture may be facilitated through this mechanism. However, this mechanism does not account for the increase in initial infection events that were only observed in the presence of ECs.

In addition to promoting infection of CD4+ T cells through potential virion capture and *trans*-infection, ECs directly engage integrins, leading to downstream changes in resting CD4+ T cells that may allow the cells to support HIV replication. Resistance of resting cells to infection results from a combination of restriction factors and low expression of HIV dependency factors [56]. While restriction factors including Murr1, SAMDH1 and Glut1 are known to be overexpressed in quiescent CD4+ T cells, these were not found to be altered in previous EC-rCD4 co-culture infection assays [21]. We investigated the possibility that co-culture of rCD4s with ECs was leading to cellular activation due to alloreactivity of CD4+ T cells to donor-mismatched ECs. Indeed, previous studies implicated MHC-II expressed by IFNγ-treated ECs in enhancement of HIV infection of resting CD4+ T cells [18,19]. However, ECs do not express MHC-II under steady state conditions and typically require IFNγ stimulation for upregulation of this pathway. We confirmed that the ECs used in our model did not express MHC-II under resting conditions or after TNFα stimulation but upregulated MHC-II after IFNγ treatment. While a low level of cellular activation of CD4+ T cells by EC co-culture was observed, the data did not support this as the main factor driving the increase in HIV infection. Specifically, of infected cells, fewer than half expressed the activation marker CD69 and fewer than 5% expressed the proliferation marker Ki67. Co-expression analysis revealed that the integrin-expressing cells identified as the primary HIV target cell were not consistently activated. Furthermore, although co-culture of ECs with rCD4s did lead to some cellular activation in the absence of HIV, only a small proportion of cells (median 3.3%) became activated as indicated by CD69 expression, compared to the large proportion of HIV-infected cells that were CD69- (median 66.2%). This is consistent with the results of previous studies that showed minimal contribution of cellular activation to enhancement of resting CD4+ T cells by ECs [20]. Nevertheless, this low-level cellular activation may promote infection of a small subset of target CD4+ T cells. Indeed, as was shown in Fig 4, there was a proportion of infected cells that expressed CD69 in the absence of other markers. Furthermore, this analysis was limited by the use of only two dynamic activation markers, CD69 and Ki67, at a single cross-sectional timepoint. A more comprehensive examination of rCD4 activation by ECs and the relative contribution of cellular activation to the enhancement of HIV infection in this system is the subject of ongoing work.

Other downstream consequences of integrin engagement may also promote establishment of productive infection. For instance, integrin signaling is a tightly controlled and dynamic process that is largely dependent on remodeling of the actin cytoskeleton. Several stages of the HIV replication cycle are similarly dependent on actin remodeling [57]. In particular, a major barrier to infection of resting CD4+ T cells occurs at the stage of nuclear import and integration of HIV DNA, which is dependent upon elements of the cytoskeleton that interact with the reverse transcriptase complex. This can be overcome by chemokine signaling, which activates the cytoskeletal factor cofilin, promoting actin rearrangement and integration of HIV DNA in resting CD4+ T cells [58]. Likewise, cytoskeletal mobilization was implicated in enhanced entry of ICAM-bearing viruses into LFA-1-expressing CD4+ T cells [59]. Similar intersecting pathways may play a role in the integrin-dependent enhancement of HIV infection in resting CD4+ T cells by ECs.

In addition to integrins, the trafficking molecule CCR6 was enriched among infected CD4+ T cells in our EC co-culture model. CCR6 is primarily expressed on Th17, Th1Th17 and Th22 CD4+ T cell subsets and binds to its ligand, CCL20, to mediate cellular migration of cells to barrier tissues such as the gut and female reproductive tract. Previous reports identified CCR6+ CD4+ T cells as preferentially infected by HIV *in vitro* [60,61] and *in vivo* [62,63]. In non-human primates, CCR6-expressing Th17 cells were shown to be the primary cellular targets of SIV and constitute the initial founder population of infected cells in the female reproductive tract [64]. These cells are enriched for HIV-dependency factors [65] and contribute to persistence of the HIV reservoir [66]. Engagement of chemokine receptors such as CCR6 by chemokines leads to integrin activation and enables resting cells to migrate into inflamed tissues. In quiescent cells, integrins exist is a closed, inactive conformation. However, upon cellular activation or chemokine signaling, inside-out signaling triggers extension of the α and β integrin subunits, thereby allowing engagement by their cognate ligands. Ligand binding leads to redistribution of integrins into clusters on the cell surface and involves cytoskeletal changes [67]. These pathways are important for HIV infection, and CCL20 signaling through CCR6 was previously shown to overcome barriers to HIV infection in resting CD4+ T cell and promote HIV latency in a manner dependent on cytoskeletal rearrangement [58]. CCL20 is produced by a variety of cell types in mucosal tissues including ECs, epithelial cells and various immune cell subsets [68]. Further characterization of chemokine production, in particular CCL20, by ECs in our co-culture model and downstream effects on integrin activation and CAM engagement by resting memory CD4+ T cells is ongoing.

The potential mechanisms described here are not necessarily mutually exclusive. It is possible that ECs contribute to infection of resting memory CD4+ T cells through *trans*-infection and through direct effects on target cells following integrin engagement. Productive infection may further be amplified in this system through integrin-dependent cell-to-cell spread between CD4+ T cells. Integrin blockade may also prevent engagement of additional interactions between ECs and resting CD4+ T cells that are dependent on cell adhesion. Furthermore, there may be additional integrin-dependent or -independent mechanisms contributing to the productive infection of resting CD4+ T cells in the presence of ECs. Consistent with this, there was a proportion of infected cells that were not characterized by high levels of integrins or activation markers. Ongoing transcriptomic studies aim to identify additional mechanisms by which ECs may promote infection of rCD4.

In summary, we demonstrate that ECs promote HIV infection of resting memory CD4+ T cells in an integrin-dependent manner. The observed enhancement of infection resulting from this interaction has clear relevance for HIV acquisition and pathogenesis. Resting memory CD4+ T cells are the primary infection targets *in vivo*, and constitute a key cellular reservoir for HIV latency. Inflammation is known to increase susceptibility to HIV acquisition and while this may be due, in part, to recruitment of activated target cells that fuel spread of infection in submucosal tissue, inflammation also has effects on the non-hematopoietic tissue stromal cells that control cell migration. Specifically, inflammatory signals upregulate cell adhesion molecules including E-selectin, ICAM and VCAM proteins on endothelial cell surfaces, leading to recruitment of resting memory and activated effector T cells into inflamed tissues. The demonstration that these proteins can enhance the HIV susceptibility of resting CD4+ T cells, that otherwise demonstrate low susceptibility to HIV infection, underscores the physiological relevance of these findings for HIV acquisition and opportunities for intervention. Treatment of inflammation or its drivers is an attractive opportunity for intervention, as it addresses the multitude of mechanisms by which inflammation may promote infection as well as additional inflammatory consequences such as tissue damage and chronic immune activation.

## Materials and Methods

### Study Subjects and Ethics Statement

Healthy blood donors were recruited locally in Winnipeg, Canada. Written consent was obtained from all study participants. The Research Ethics Board at the University of Manitoba approved the study protocols. For some experiments, whole blood was purchased from StemCell Technologies.

### Cell Isolation and Culture

Peripheral blood mononuclear cells (PBMC) were isolated from whole blood by density gradient centrifugation. Resting CD4+ T cells (rCD4s) were isolated from PBMC by magnetic negative selection using the EasySep Human Resting CD4+ T cell Isolation Kit (StemCell Technologies). In some experiments, bulk CD4+ T cells were isolated using the EasySep CD4+ T cell Enrichment Kit (StemCell Technologies). Isolated CD4+ T cells were maintained in RPMI medium supplemented with 10% fetal bovine serum and 1% penicillin and streptomycin (R10 medium). Purity of isolated cells was assessed by flow cytometry.

Human umbilical vein endothelial cells (ECs) were obtained from Lonza and cultured in EGM-2 complete culture medium (Lonza). ECs were seeded at 5000 cells/cm^2^ and cultured to at least 70% confluence before co-culture with CD4+ T cells.

### EC-rCD4 co-cultures and infection assays

ECs were cultured to at least 70% confluence. Where indicated, ECs were treated with 100U/ml TNFα (R&D Systems) overnight to maximize expression of cell adhesion molecules prior to co-culture. At the time of co-culture, 300,000 rCD4s were added per well to EC monolayers in a 48 well plate. Where indicated, rCD4 cells were pre-incubated with blocking antibodies (10μg/ml anti-αL clone TS1/22 + 2.5 μg/ml anti-α4 clone 2B4) or isotype control antibody (12.5μg/ml mouse IgG1κ clone 11711) for 30 minutes before being added to ECs then the blocking antibodies were maintained in the co-culture throughout the experiment. Co-cultures were maintained in RPMI medium supplemented with 10% fetal bovine serum and 1% penicillin and streptomycin. After 24 hours, co-cultures were exposed to HIV_IIIB_ at a MOI of 1.0 pfu/rCD4 target cell. After 4 hours, co-cultures were centrifuged to remove input virus then maintained in R10 medium with 10U/ml recombinant IL-2 for three days. On day 3 post-infection, rCD4s were collected and analyzed by flow cytometry. Supernatants were collected from culture wells, treated with 1% Triton X-100 (Sigma Aldrich) and stored at -80°C.

### Flow Cytometry

Cells were washed then incubated with a cocktail of cell surface antibodies at 4°C for 30 minutes. Cells were then washed and either fixed with 1% paraformaldehyde for 30 minutes or further stained for intracellular antigens. Intracellular staining of HIV-p24 and Ki67 was performed using the eBioscience Foxp3 / Transcription Factor Staining Buffer set (ThermoFisher Scientific). A list of flow cytometry panels and antibodies used is provided in S3 Table. Samples were acquired on a BD LSRII using FACSDiva software (BD Biosciences) and data were analyzed using FlowJo software. Positive fluorescence signals were established using fluorescence minus one (FMO) controls. Visualization of co-expression analysis was performed using SPICE software [69].

### HIV-p24 ELISAs

Culture supernatants were diluted 20-fold with PBS + 1% BSA + 0.2% Triton X-100 then analyzed for HIV-p24 protein using the HIV-1 Gag p24 DuoSet ELISA Development System (R&D Systems) with the R&D TMB substrate reagent pack for detection. Reaction was stopped using 2N H2SO4 then optical density at 450nm with wavelength correction at 570nm was acquired using a Synergy H1 plate reader.

### Static Adhesion Assays

Titration of blocking antibodies was performed by using static adhesion assays in which activated CD4+ T cells were allowed to adhere to either recombinant human ICAM/Fc or VCAM/Fc chimeras in the presence of blocking antibodies specific for integrin α4 (clone 2B4, R&D Systems) or integrin αL (clone TS1/22, ThermoFisher) or mouse IgG1κ isotype control (clone 11711, R&D Systems). Wells were coated with 5μg/ml of either rhICAM/Fc chimera (R&D Systems) or rhVCAM/Fc chimera (R&D Systems) overnight at 4°C. Plates were washed then blocked with PBS containing 2% Goat Serum and 0.01% Tween-20 for 2 hours at 37°C. Purified CD4+ T cells were counted and labelled with 2 μM Cell Tracker Green CMFDA dye (ThermoFisher) for 20 minutes at 37°C. Cells were then stimulated overnight with 100ng/ml PMA and 1μM ionomycin. Activated CD4+ T cells were pre-incubated with blocking antibodies for 30 minutes at 37°C. Cells were transferred to rhCAM/Fc-coated plates (5 x 10^4^/well) and allowed to adhere in the presence or absence of blocking antibody or isotype control for 30 minutes at 37°C. Wells were gently mixed and supernatant was aspirated to remove non-adherent cells. Wells were washed gently twice with PBS then adherent cells were visualized using GFP channel on an EVOS fluorescent microscope. Digital images were captured under the same magnification and intensification. Image processing was restricted to adjustment of levels using Adobe Photoshop and all images acquired under the same conditions were treated identically. Cell-tracker green labelled cells were counted using Cell Profiler software. A minimum of 3 fields was counted per well in a 96 well plate.

To confirm effect of blocking in the EC-rCD4 co-culture experiments, the static adhesion assay was performed using EC monolayers instead of rhCAM/Fc chimeras. EC were grown to at least 70% confluence then media was aspirated and wells were rinsed with PBS. Purified rCD4 were labelled with Cell Tracker Green CMFDA dye as above. Labelled rCD4 were pre-incubated with blocking antibodies (10μg/ml anti-αL clone TS1/22 + 2.5 μg/ml anti-α4 clone 2B4) or isotype control antibody (12.5μg/ml mouse IgG1κ clone 11711) for 30 minutes where indicated, then added to EC monolayers and allowed to adhere in the presence of antibody for 30 minutes at 37°C. As above, wells were mixed gently then non-adhered cells were aspirated and wells were washed twice before visualization and image capture on an EVOS microscope.

### Statistical Analysis

Continuous variables in matched samples were analyzed using non-parametric paired Friedman ANOVA followed by Dunn’s multiple comparisons test when analyzing three or more groups. For the analysis of ELISA data in Fig 2, non-parametric unpaired Kruskal-Wallis ANOVA had to be used due to lack of supernatant available for analysis of the HUVEC condition for two participants. For comparison of two matched samples, the non-parametric Wilcoxon matched-pairs signed rank test was used. Differences were considered to be statistically significant if p < 0.05. All statistical analyses were performed using GraphPad Prism software, version 8 except for comparisons of phenotypes in the co-expression analyses, which were done within SPICE software [69]. In those analyses, asterisks indicate statistically-significant differences meeting the threshold for Bonferroni correction.

## Acknowledgements

The authors would like to thank the volunteers that donated samples for this study. We would also like to thank Linda Ares, Xuefen Yang and Tomasz Bielawny for assistance with sample coordination.

## Supporting Information

**S1 Figure.**
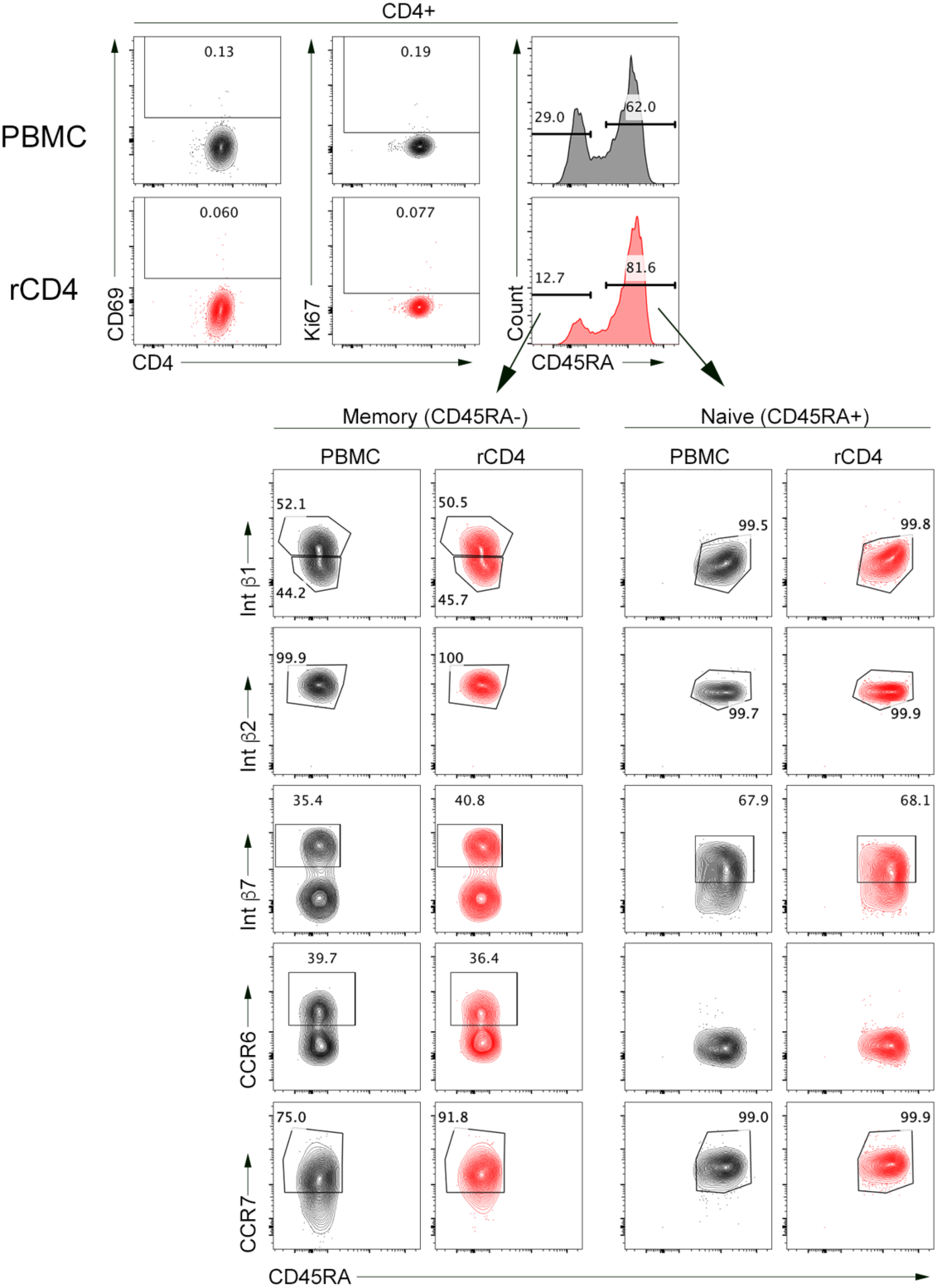
PBMC (grey) and rCD4 (red) were assessed for markers of activation (CD69, Ki67), memory (CD45RA, CCR7), integrins VLA-4 (β1), LFA-1 (β2) and α4β7 (β7) and CCR6. Quantification and statistical comparisons are shown in Table 1.

**S2 Figure.**
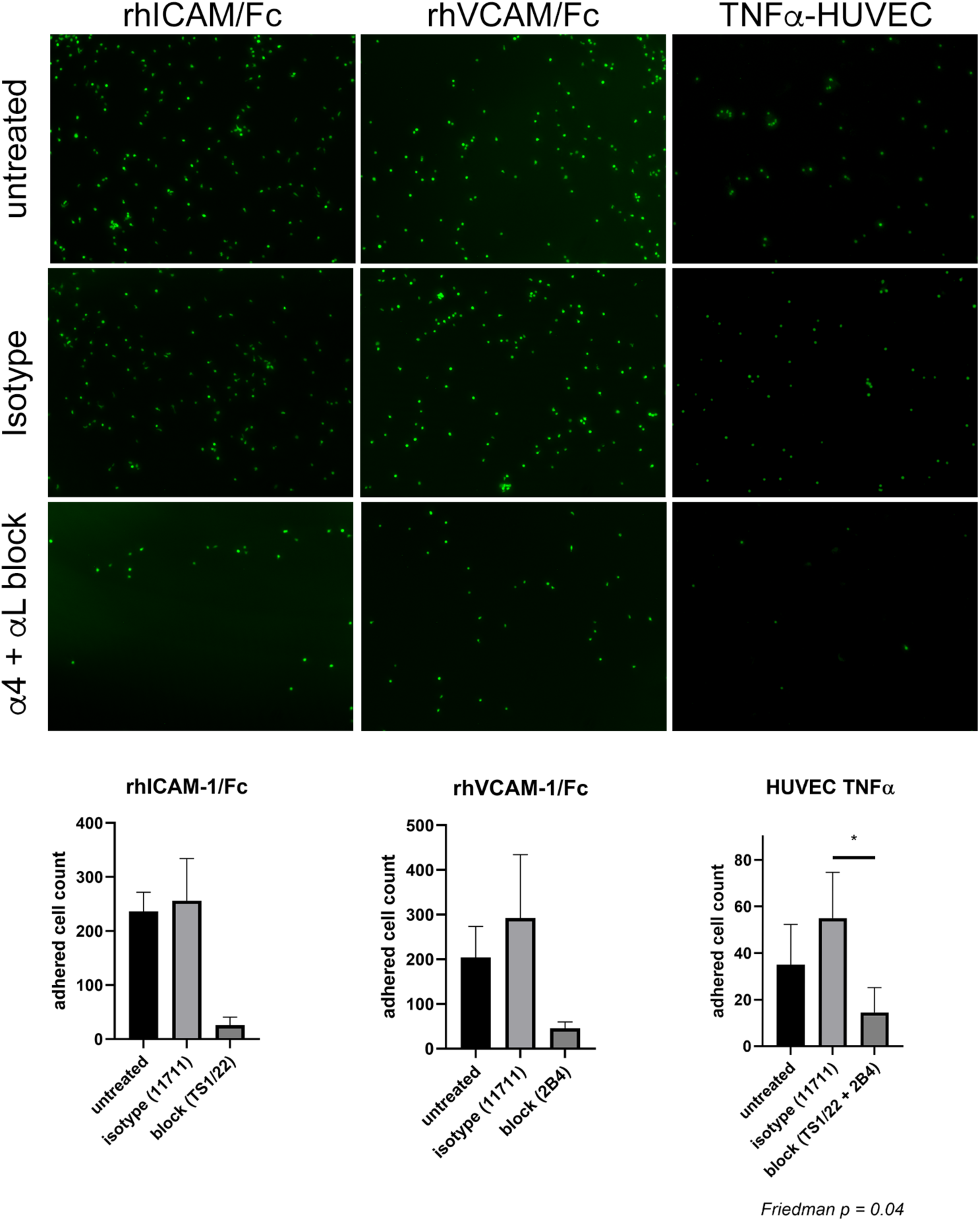
Integrin-specific blocking antibodies prevent cell adhesion. CD4+ T cells were labelled with Cell Tracker Green and allowed to adhere to either rhICAM-1/Fc, rhVCAM-1/Fc or a monolayer of TNFα-treated HUVECs then non-adhered cells were removed and adhered cells were visualized using fluorescence microscopy. Antibodies specific for LFA-1 (αL, clone TS1/22) and VLA-4 (α4, clone 2B4) inhibited adhesion to rhICAM-1/Fc and rhVCAM-1 Fc, respectively. The combination of blocking antibodies prevented adhesion to HUVEC monolayers. A mouse IgG1 isotype control antibody (clone 11711) had no effect.

**S3 Figure.**
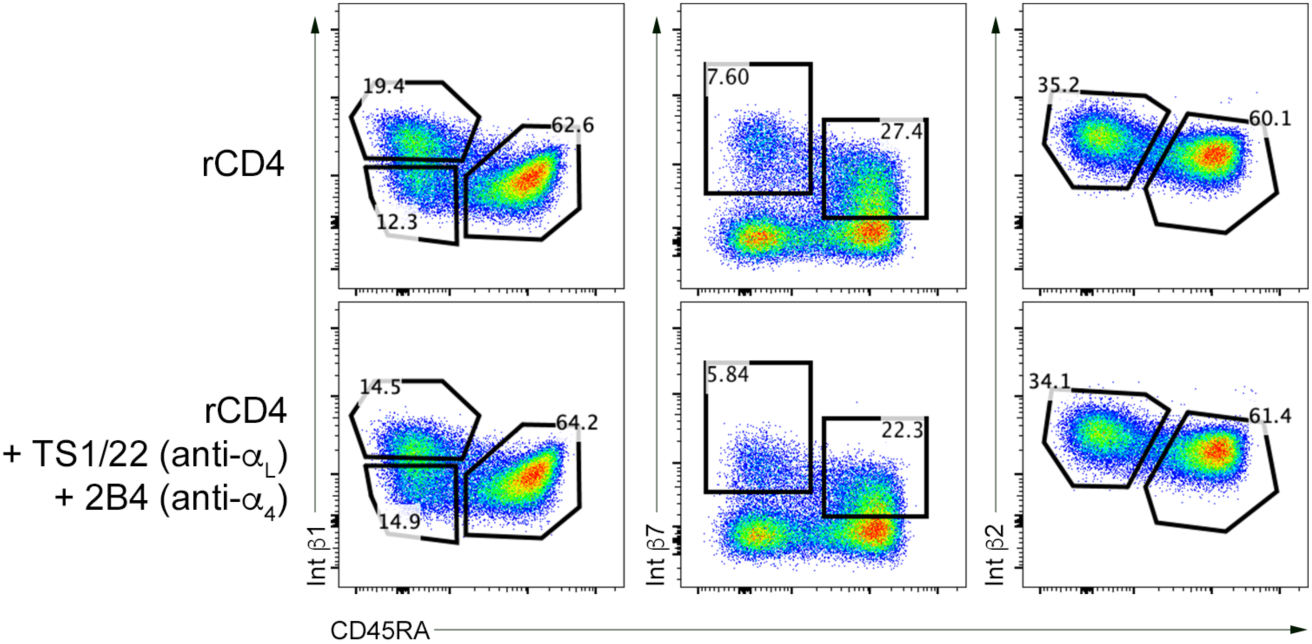
Anti-α4 antibody blocks detection of β1 and β7 integrin chains. rCD4 cells were cultured alone or in the presence of anti-αL (clone TS1/22) and anti-α4 (clone 2B4) for 30 minutes before staining for integrin β chains β1, β2 and β7. Blocking interfered with detection of integrin β1 (β chain of VLA-4) and β7 (β chain of α4β7) but not β2 (β chain of LFA-1).

**S4 Figure.**
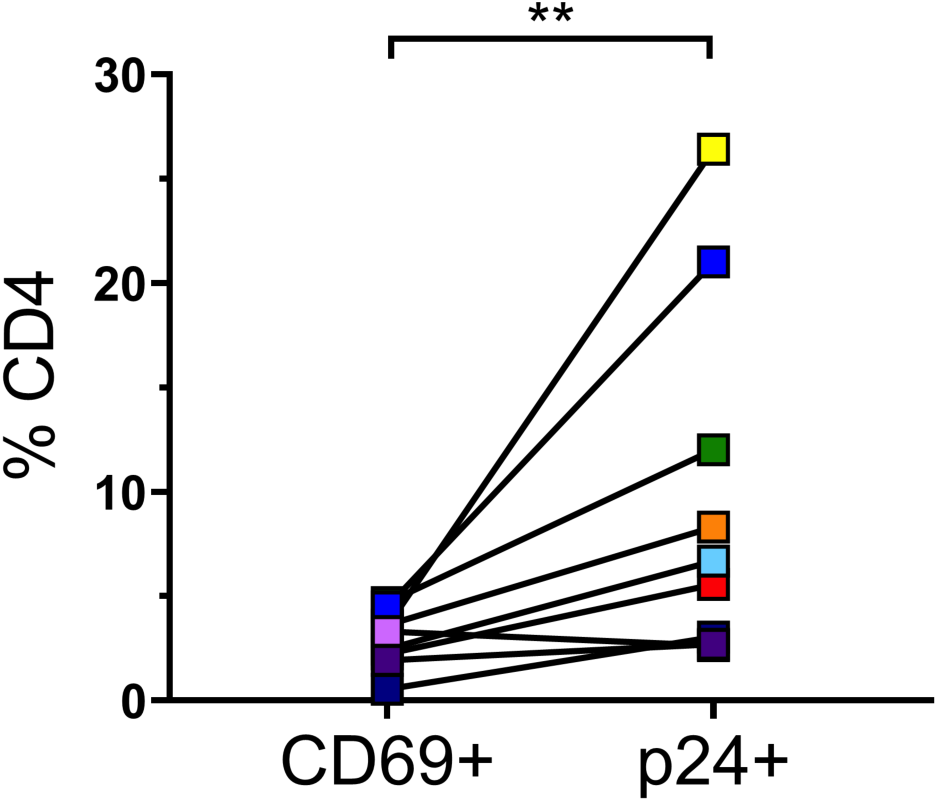
Comparison of rCD4 cells that became activated after co-culture with ECs (CD69+) and cells that became infected after co-culture with ECs and exposure to HIV_IIIB_ (p24+). Statistical comparison was performed by Wilcoxon matched-pairs signed-rank test (** p < 0.01).

**S1 Table.** Comparison of phenotypes between HIV-p24- and HIV-p24+ CD4+ T cells in co-cultures with TNFα-HUVEC

| Parameter <sup>a</sup> | HIV-p24- | HIV-p24+ | p-value <sup>b</sup> |
| --- | --- | --- | --- |
| CD45RA- CCR7+ (%) | 7.1 (4.4-10.3) | 52.5 (38.1-71.0) | **0.004** |
| CD45RA+ CCR7+ (%) | 87.2 (81.4-92.3) | 33.2 (19.9-47.3) | **0.004** |
| $\beta 1^{\text{hi}}$ CD45RA- (%) | 5.0 (2.1-11.1) | 43.3 (32.2-57.3) | **0.004** |
| $\beta 1$ MFI | 2709 (2500-3046) | 3460 (2946-3561) | **0.004** |
| $\beta 1^{\text{lo}}$ CD45RA- (%) | 4.2 (2.6-6.6) | 14.6 (11.6-18.5) | **0.004** |
| $\beta 1$ MFI | 924.5 (895.0-945.0) | 1133 (1091-1163) | **0.004** |
| $\beta 1^{\text{lo}}$ CD45RA+ (%) | 88.4 (83.3-93.4) | 32.8 (16.7-42.2) | **0.004** |
| $\beta 1$ MFI | 833.0 (786.5-953.0) | 988.0 (875.5-1160) | **0.004** |
| $\beta 2^{\text{hi}}$ CD45RA- (%) | 8.9 (4.6-12.4) | 63.8 (49.1-77.0) | **0.004** |
| $\beta 2$ MFI | 2500 (2223-2709) | 11247 (4008-14093) | **0.004** |
| $\beta 2^+$ CD45RA+ (%) | 90.5 (85.8-94.3) | 35.5 (16.4-44.5) | **0.004** |
| $\beta 2$ MFI | 4516 (220.0-5241) | 5506 (1882-6399) | **0.004** |
| $\beta 7^{\text{hi}}$ CD45RA- (%) | 2.6 (1.2-3.5) | 16.3 (11.3-18.1) | **0.004** |
| $\beta 7$ MFI | 4220 (2069-4903) | 4319 (2063-4530) | 0.250 |
| $\beta 7^{\text{int}}$ CD45RA+ (%) | 53.6 (42.4-58.6) | 22.3 (12.0-28.9) | **0.004** |
| $\beta 7$ MFI | 914.0 (415.0-1251) | 1070 (443.5-1408) | **0.020** |
| CCR6 (%) | 1.07 (0.28-2.6) | 34.3 (13.6-45.1) | **0.004** |
| CCR6 MFI | 1312 (1196-1390) | 2099 (1839-2225) | **0.004** |
| CD69 (%) | 1.1 (0.42-2.0) | 33.8 (22.9-37.0) | **0.004** |
| Ki67 (%) | 0.04 (0.02-0.06) | 2.7 (2.0-3.0) | **0.004** |

<sup>a</sup> expression values listed as median of % positive (+) cells within HIV-p24+ or HIV-p24- parent population or median fluorescence intensity (MFI) of indicated cell population, with interquartile range shown in parentheses
<sup>b</sup> p-values calculated using non-parametric Wilcoxon matched pairs test. Parameters considered statistically significant (p<0.05) are highlighted in bold text

**S2 Table.** Comparison of phenotypes between *ex vivo* isolated rCD4 and rCD4 after culture for 3 days alone or with TNFα-treated HUVEC

| Parameter <sup>a</sup> | *ex vivo* | 4 days culture |  | p-value <sup>b</sup> |
| --- | --- | --- | --- | --- |
| | rCD4 | rCD4 alone | HUVEC-TNFα | |
| CD4+ T cell populations |  |  |  |  |
| CD69 (%CD4+) | 0.06 (0.03-0.42) | 0.05 (0.03-1.40) | 3.30 (2.01-4.05) | **0.0035** |
| Ki67 (%CD4+) | 0.09 (0.05-0.45) | 0.04 (0.02-0.09) | 1.15 (0.61-0.57) | **<0.0001** |
| CD45RA (%CD4+) | 65.5 (59.1-75.7) | 66.8 (63.7-77.5) | 77.5 (68.5-84.9) | **0.0002** |
| Naïve CD4+ T cell populations |  |  |  |  |
| $\beta 1^{\text{lo}}$ (%CD45RA+) | 99.6 (99.4-99.8) | 99.2 (98.6-99.5) | 99.1 (98.5-99.7) | 0.064 |
| $\beta 1$ MFI | 853.0 (742.5-895.5) | 735.0 (615.0-910.5) | 793.0 (756.0-911.5) | 0.569 |
| $\beta 2^+$ (%CD45RA+) | 99.9 (99.8-100.0) | 99.8 (99.7-100.0) | 99.7 (99.4-100) | 0.123 |
| $\beta 2$ MFI | 5552 (1778-5858) | 3007 (1251-3399) | 4334 (1122-3399) | **0.0002** |
| $\beta 7^{\text{int}}$ (%CD45RA+) | 54.3 (43.3-61.7) | 65.8 (57.4-77.9) | 71.0 (54.7-77.3) | **0.0013** |
| $\beta 7$ MFI | 944.0 (452.5-1058) | 1081 (430.0-1483) | 1012 (493.5-1390) | 0.154 |
| CCR7 (%CD45RA+) | 99.3 (98.8-99.9) | 99.2 (98.8-99.6) | 99.3 (98.7-99.5) | 0.814 |
| Memory CD4+ T cell populations |  |  |  |  |
| $\beta 1^{\text{hi}}$ (%CD45RA-) | 52.9 (48.6-58.3) | 47.8 (41.5-70.1) | 58.4 (43.4-67.2) | 0.814 |
| $\beta 1$ MFI | 2021 (1912-2596) | 2899 (2733-3511) | 2766 (2536-3728) | **0.0003** |
| $\beta 1^{\text{lo}}$ (%CD45RA-) | 38.7 (36.4-48.3) | 48.9 (24.2-54.8) | 36.9 (27.3-53.4) | 0.814 |
| $\beta 1$ MFI | 720.0 (628.5-853.0) | 905.0 (842.5-949.0) | 807.0 (787.5-998.0) | **0.0013** |
| $\beta 2^{\text{hi}}$ (%CD45RA-) | 99.8 (99.8-100.0) | 99.8 (98.8-100.0) | 99.9 (99.8-100) | 0.329 |
| $\beta 2$ MFI | 8053 (2981-9328) | 5660 (2965-6491) | 9743 (2234-13046) | 0.278 |
| $\beta 7^{\text{hi}}$ (%CD45RA-) | 25.6 (20.9-31.6) | 29.2 (20.9-40.6) | 29.9 (26.2-37.4) | **0.0035** |
| $\beta 7^{\text{hi}}$ MFI | 3388 (1971-3733) | 6892 (2560-7272) | 5297 (2887-6013) | **<0.0001** |
| CCR6 (%CD45RA-) | 40.2 (31.2-49.2) | 29.5 (23.8-38.5) | 29.0 (18.8-34.2) | **0.0007** |
| CCR6 MFI | 2433 (2230-2824) | 2193 (2088-2464) | 2728 (1879-3009) | 0.741 |
| CCR7 (%CD45RA-) | 87.7 (84.8-91.7) | 89.5 (79.9-92.6) | 91.2 (89.6-93.2) | 0.057 |

<sup>a</sup> expression values listed as median of % positive (+) cells in indicated parent population or median fluorescence intensity (MFI), with interquartile range shown in parentheses
<sup>b</sup> p-values calculated using non-parametric Friedman test. Parameters considered statistically significant (p<0.05) are highlighted in bold text

**S3 Table.**
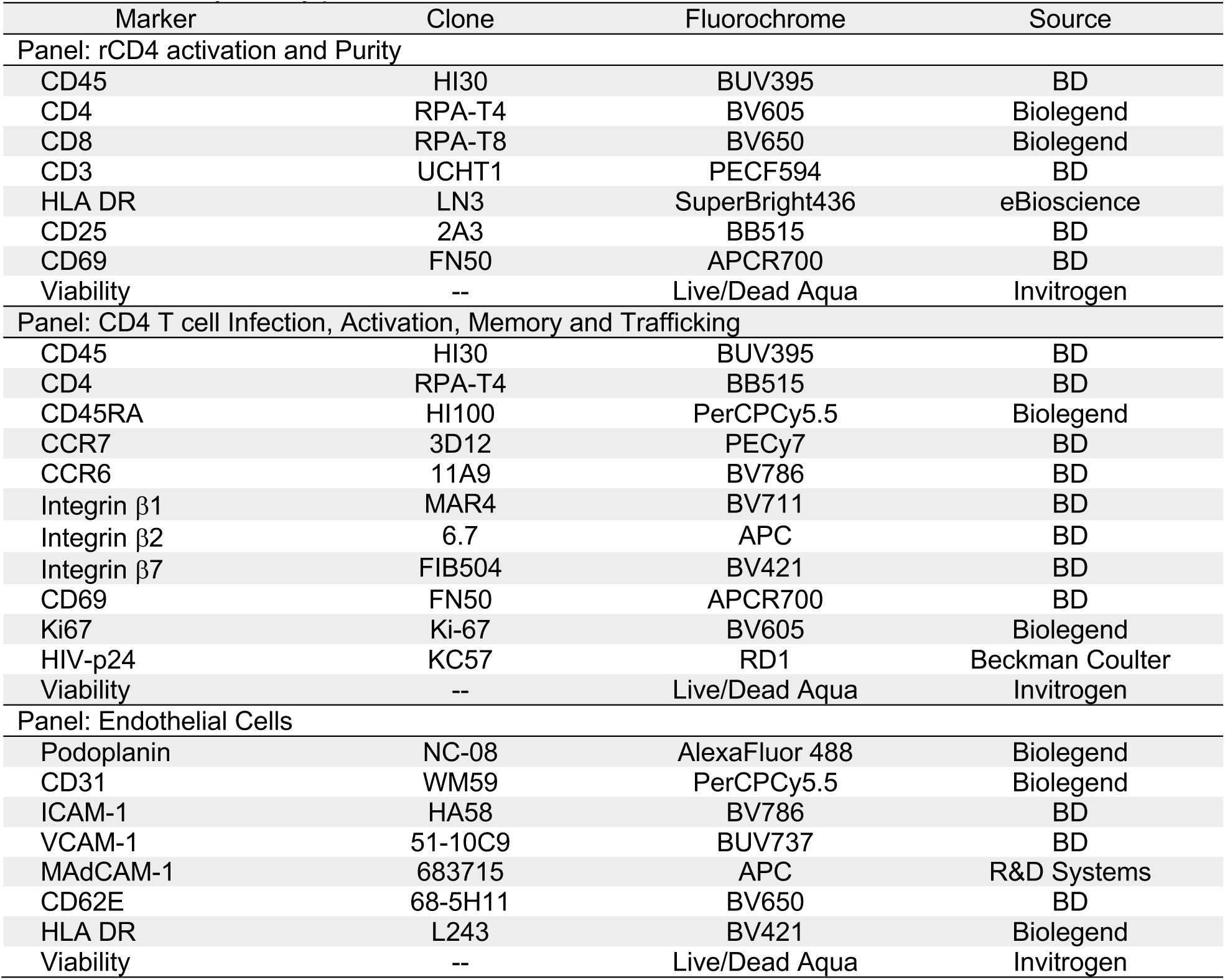
Flow cytometry panels

## References

1. Masson L, Passmore J-AS, Liebenberg LJ, Werner L, Baxter C, Arnold KB, et al. Genital inflammation and the risk of HIV acquisition in women. Clin Infect Dis. 2015;61: 260–269. doi:10.1093/cid/civ298

2. McKinnon LR, Liebenberg LJ, Yende-Zuma N, Archary D, Ngcapu S, Sivro A, et al. Genital inflammation undermines the effectiveness of tenofovir gel in preventing HIV acquisition in women. Nature Publishing Group. Nature Publishing Group; 2018;24: 491–496. doi:10.1038/nm.4506

3. Nazli A, Chan O, Dobson-Belaire WN, Ouellet M, Tremblay MJ, Gray-Owen SD, et al. Exposure to HIV-1 directly impairs mucosal epithelial barrier integrity allowing microbial translocation. PLoS Pathog. 2010;6: e1000852. doi:10.1371/journal.ppat.1000852

4. Kaul R, Rebbapragada A, Hirbod T, Wachihi C, Ball TB, Plummer FA, et al. Genital levels of soluble immune factors with anti-HIV activity may correlate with increased HIV susceptibility. AIDS. 2008;22: 2049–2051. doi:10.1097/QAD.0b013e328311ac65

5. Spina CA, Guatelli JC, Richman DD. Establishment of a stable, inducible form of human immunodeficiency virus type 1 DNA in quiescent CD4 lymphocytes in vitro. J Virol. American Society for Microbiology; 1995;69: 2977–2988.

6. Korin YD, Zack JA. Progression to the G1b phase of the cell cycle is required for completion of human immunodeficiency virus type 1 reverse transcription in T cells. J Virol. 1998;72: 3161–3168.

7. Stevenson M, Stanwick TL, Dempsey MP, Lamonica CA. HIV-1 replication is controlled at the level of T cell activation and proviral integration. The EMBO Journal. European Molecular Biology Organization; 1990;9: 1551–1560.

8. Zack JA, Arrigo SJ, Weitsman SR, Go AS, Haislip A, Chen IS. HIV-1 entry into quiescent primary lymphocytes: molecular analysis reveals a labile, latent viral structure. Cell. 1990;61: 213–222.

9. Zack JA, Haislip AM, Krogstad P, Chen IS. Incompletely reverse-transcribed human immunodeficiency virus type 1 genomes in quiescent cells can function as intermediates in the retroviral life cycle. J Virol. 1992;66: 1717–1725. doi:10.1016/0169-6009(92)92142-d

10. Zhou Y, Zhang H, Siliciano JD, Siliciano RF. Kinetics of human immunodeficiency virus type 1 decay following entry into resting CD4+ T cells. J Virol. 2005;79: 2199–2210. doi:10.1128/JVI.79.4.2199-2210.2005

11. Zhang ZQ, Zhang ZQ, Schuler T, Schuler T, Zupancic M, Zupancic M, et al. Sexual transmission and propagation of SIV and HIV in resting and activated CD4+ T cells. Science. American Association for the Advancement of Science; 1999;286: 1353.

12. Li Q, Duan L, Estes JD, Ma Z-M, Rourke T, Wang Y, et al. Peak SIV replication in resting memory CD4+ T cells depletes gut lamina propria CD4+ T cells. Nature. 2005;434: 1148–1152. doi:10.1038/nature03513

13. Kinter A, Moorthy A, Jackson R, Fauci AS. Productive HIV infection of resting CD4+ T cells: role of lymphoid tissue microenvironment and effect of immunomodulating agents. AIDS Res Hum Retroviruses. 2003;19: 847–856. doi:10.1089/088922203322493012

14. Eckstein DA, Penn ML, Korin YD, Scripture-Adams DD, Zack JA, Kreisberg JF, et al. HIV-1 actively replicates in naive CD4(+) T cells residing within human lymphoid tissues. Immunity. 2001;15: 671–682.

15. Nishimura Y, Brown CR, Mattapallil JJ, Igarashi T, Buckler-White A, Lafont BAP, et al. Resting naive CD4+ T cells are massively infected and eliminated by X4-tropic simianhuman immunodeficiency viruses in macaques. Proceedings of the National Academy of Sciences. 2005;102: 8000–8005. doi:10.1073/pnas.0503233102

16. Ostrowski MA, Chun TW, Justement SJ, Motola I, Spinelli MA, Adelsberger J, et al. Both memory and CD45RA+/CD62L+ naive CD4(+) T cells are infected in human immunodeficiency virus type 1-infected individuals. J Virol. 1999;73: 6430–6435.

17. Cary DC, Fujinaga K, Peterlin BM. Molecular mechanisms of HIV latency. J Clin Invest. 2016;126: 448–454. doi:10.1172/JCI80565

18. Choi J, Walker J, Talbert-Slagle K, Wright P, Pober JS, Alexander L. Endothelial cells promote human immunodeficiency virus replication in nondividing memory T cells via Nef-, Vpr-, and T-cell receptor-dependent activation of NFAT. J Virol. 2005;79: 11194–11204. doi:10.1128/JVI.79.17.11194-11204.2005

19. Choi J, Walker J, Boichuk S, Kirkiles-Smith N, Torpey N, Pober JS, et al. Human endothelial cells enhance human immunodeficiency virus type 1 replication in CD4+ T cells in a Nef-dependent manner in vitro and in vivo. J Virol. 2005;79: 264–276. doi:10.1128/JVI.79.1.264-276.2005

20. Shen A, Baker JJ, Scott GL, Davis YP, Ho Y-Y, Siliciano RF. Endothelial cell stimulation overcomes restriction and promotes productive and latent HIV-1 infection of resting CD4+ T cells. J Virol. 2013;87: 9768–9779. doi:10.1128/JVI.01478-13

21. Morris JH, Nguyen T, Nwadike A, Geels ML, Kamp DL, Kim BR, et al. Soluble Factors Secreted by Endothelial Cells Allow for Productive and Latent HIV-1 Infection in Resting CD4(+) T Cells. AIDS Res Hum Retroviruses. 2016. doi:10.1089/AID.2016.0058

22. Schilthuis M, Verkaik S, Walhof M, Philipose A, Harlow O, Kamp D, et al. Lymphatic endothelial cells promote productive and latent HIV infection in resting CD4+ T cells. Virol J. BioMed Central; 2018;15: 152. doi:10.1186/s12985-018-1068-6

23. Thibault S, Tardif MR, Gilbert C, Tremblay MJ. Virus-associated host CD62L increases attachment of human immunodeficiency virus type 1 to endothelial cells and enhances trans infection of CD4+ T lymphocytes. Journal of General Virology. 2007;88: 2568–2573. doi:10.1099/vir.0.83032-0

24. Bobardt MD, Saphire ACS, Hung H-C, Yu X, Van der Schueren B, Zhang Z, et al. Syndecan captures, protects, and transmits HIV to T lymphocytes. Immunity. 2003;18: 27–39. doi:10.1016/s1074-7613(02)00504-6

25. Bounou S, Leclerc JE, Tremblay MJ. Presence of Host ICAM-1 in Laboratory and Clinical Strains of Human Immunodeficiency Virus Type 1 Increases Virus Infectivity and CD4+-T-Cell Depletion in Human Lymphoid Tissue, a Major Site of Replication In Vivo. J Virol. 2002;76: 1004–1014. doi:10.1128/JVI.76.3.1004-1014.2002

26. Fortin JF, Cantin R, Tremblay MJ. T cells expressing activated LFA-1 are more susceptible to infection with human immunodeficiency virus type 1 particles bearing host-encoded ICAM-1. J Virol. 1998;72: 2105–2112.

27. Tardif MR, Tremblay MJ. LFA-1 Is a Key Determinant for Preferential Infection of Memory CD4+ T Cells by Human Immunodeficiency Virus Type 1. J Virol. 2005;79: 13714–13724. doi:10.1128/JVI.79.21.13714-13724.2005

28. Piguet V, Sattentau Q. Dangerous liaisons at the virological synapse. Journal of Clinical Investigation. 2004;114: 605–610. doi:10.1172/JCI22812

29. Bracq L, Xie M, Benichou S, Bouchet J. Mechanisms for Cell-to-Cell Transmission of HIV-1. Front Immunol. Frontiers; 2018;9: 260. doi:10.3389/fimmu.2018.00260

30. Tardif MR, Gilbert C, Thibault S, Fortin JF, Tremblay MJ. LFA-1 Antagonists as Agents Limiting Human Immunodeficiency Virus Type 1 Infection and Transmission and Potentiating the Effect of the Fusion Inhibitor T-20. Antimicrob Agents Chemother. 2009;53: 4656–4666. doi:10.1128/AAC.00117-09

31. Joag VR, McKinnon LR, Liu J, Kidane ST, Yudin MH, Nyanga B, et al. Identification of preferential CD4+ T-cell targets for HIV infection in the cervix. Mucosal Immunology. Nature Publishing Group; 2016;9: 1–12. doi:10.1038/mi.2015.28

32. Kader M, Wang X, Piatak M, Lifson J, Roederer M, Veazey R, et al. Alpha4(+)beta7(hi)CD4(+) memory T cells harbor most Th-17 cells and are preferentially infected during acute SIV infection. Mucosal Immunology. 2009;2: 439–449. doi:10.1038/mi.2009.90

33. Wang X, Xu H, Gill AF, Pahar B, Kempf D, Rasmussen T, et al. Monitoring alpha4beta7 integrin expression on circulating CD4+ T cells as a surrogate marker for tracking intestinal CD4+ T-cell loss in SIV infection. Mucosal Immunology. 2009;2: 518–526. doi:10.1038/mi.2009.104

34. Cicala C, Martinelli E, McNally JP, Goode DJ, Gopaul R, Hiatt J, et al. The integrin alpha4beta7 forms a complex with cell-surface CD4 and defines a T-cell subset that is highly susceptible to infection by HIV-1. Proc Natl Acad Sci USA. 2009;106: 20877–20882. doi:10.1073/pnas.0911796106

35. McKinnon LR, Nyanga B, Chege D, Izulla P, Kimani M, Huibner S, et al. Characterization of a Human Cervical CD4+ T Cell Subset Coexpressing Multiple Markers of HIV Susceptibility. The Journal of Immunology. 2011;187: 6032–6042. doi:10.4049/jimmunol.1101836

36. Lu X, Li Z, Li Q, Jiao Y, Ji Y, Zhang H, et al. Preferential loss of gut-homing α4β7 CD4(+) T cells and their circulating functional subsets in acute HIV-1 infection. Cell Mol Immunol. 2016;13: 776–784. doi:10.1038/cmi.2015.60

37. Martinelli E, Veglia F, Goode D, Guerra-Perez N, Aravantinou M, Arthos J, et al. The frequency of α4β+ (high) memory CD4+ T cells correlates with susceptibility to rectal simian immunodeficiency virus infection. J Acquir Immune Defic Syndr. 2013;64: 325–331. doi:10.1097/QAI.0b013e31829f6e1a

38. Nawaz F, Goes LR, Ray JC, Olowojesiku R, Sajani A, Ansari AA, et al. MAdCAM costimulation through Integrin-Î±4Î^2^7 promotes HIV replication. Mucosal Immunology. Springer US; 2018;: 1–10. doi:10.1038/s41385-018-0044-1

39. Byrareddy SN, Kallam B, Arthos J, Cicala C, Nawaz F, Hiatt J, et al. Targeting α4β7 integrin reduces mucosal transmission of simian immunodeficiency virus and protects gut-associated lymphoid tissue from infection. Nat Med. 2014;20: 1397–1400. doi:10.1038/nm.3715

40. Byrareddy SN, Arthos J, Cicala C, Villinger F, Ortiz KT, Little D, et al. Sustained virologic control in SIV+ macaques after antiretroviral and α4β7 antibody therapy. Science. 2016;354: 197–202. doi:10.1126/science.aag1276

41. Iwamoto N, Mason RD, Song K, Gorman J, Welles HC, Arthos J, et al. Blocking α4β7 integrin binding to SIV does not improve virologic control. Science. 2019;365: 1033–1036. doi:10.1126/science.aaw7765

42. Abbink P, Mercado NB, Nkolola JP, Peterson RL, Tuyishime H, McMahan K, et al. Lack of therapeutic efficacy of an antibody to α4β7 in SIVmac251-infected rhesus macaques. Science. 2019;365: 1029–1033. doi:10.1126/science.aaw8562

43. Di Mascio M, Lifson JD, Srinivasula S, Kim I, DeGrange P, Keele BF, et al. Evaluation of an antibody to α4β7 in the control of SIVmac239-nef-stop infection. Science. American Association for the Advancement of Science; 2019;365: 1025–1029. doi:10.1126/science.aav6695

44. Sneller MC, Clarridge KE, Seamon C, Shi V, Zorawski MD, Justement JS, et al. An open-label phase 1 clinical trial of the anti-α4β7 monoclonal antibody vedolizumab in HIV- infected individuals. Science Translational Medicine. 2019;11: eaax3447. doi:10.1126/scitranslmed.aax3447

45. Uotila LM, Jahan F, Soto Hinojosa L, Melandri E, Grönholm M, Gahmberg CG. Specific phosphorylations transmit signals from leukocyte β2 to β1 integrins and regulate adhesion. Journal of Biological Chemistry. 2014;289: 32230–32242. doi:10.1074/jbc.M114.588111

46. Grönholm M, Jahan F, Bryushkova EA, Madhavan S, Aglialoro F, Soto Hinojosa L, et al. LFA-1 integrin antibodies inhibit leukocyte α4β1-mediated adhesion by intracellular signaling. Blood. 2016;128: 1270–1281. doi:10.1182/blood-2016-03-705160

47. Tremblay MJ, Fortin JF, Cantin R. The acquisition of host-encoded proteins by nascent HIV-1. Immunol Today. 1998;19: 346–351. doi:10.1016/s0167-5699(98)01286-9

48. Martin G, Tremblay MJ. HLA-DR, ICAM-1, CD40, CD40L, and CD86 are incorporated to a similar degree into clinical human immunodeficiency virus type 1 variants expanded in natural reservoirs such as peripheral blood mononuclear cells and human lymphoid tissue cultured ex vivo. Clinical Immunology. 2004;111: 275–285. doi:10.1016/j.clim.2004.02.004

49. Guzzo C, Ichikawa D, Park C, Phillips D, Liu Q, Zhang P, et al. Virion incorporation of integrin α4β7 facilitates HIV-1 infection and intestinal homing. Science Immunology. Science Immunology; 2017;2: eaam7341. doi:10.1126/sciimmunol.aam7341

50. Liao Z, Roos JW, Hildreth JE. Increased infectivity of HIV type 1 particles bound to cell surface and solid-phase ICAM-1 and VCAM-1 through acquired adhesion molecules LFA- 1 and VLA-4. AIDS Res Hum Retroviruses. Mary Ann Liebert, Inc; 2000;16: 355–366. doi:10.1089/088922200309232

51. Asin SN, Fanger MW, Wildt-Perinic D, Ware PL, Wira CR, Howell AL. Transmission of HIV-1 by primary human uterine epithelial cells and stromal fibroblasts. J Infect Dis. 2004;190: 236–245. doi:10.1086/421910

52. Meng G, Wei X, Wu X, Sellers MT, Decker JM, Moldoveanu Z, et al. Primary intestinal epithelial cells selectively transfer R5 HIV-1 to CCR5+ cells. Nat Med. 2002;8: 150–156. doi:10.1038/nm0202-150

53. Neidleman JA, Chen JC, Kohgadai N, Müller JA, Laustsen A, Thavachelvam K, et al. Mucosal stromal fibroblasts markedly enhance HIV infection of CD4+ T cells. PLoS Pathog. 2017;13: e1006163. doi:10.1371/journal.ppat.1006163

54. Murakami T, Kim J, Li Y, Green GE, Shikanov A, Ono A. Secondary lymphoid organ fibroblastic reticular cells mediate trans-infection of HIV-1 via CD44- hyaluronan interactions. Nat Comms. Springer US; 2018;: 1–14. doi:10.1038/s41467-018-04846-w

55. Jolly C, Mitar I, Sattentau QJ. Adhesion molecule interactions facilitate human immunodeficiency virus type 1-induced virological synapse formation between T cells. J Virol. 2007;81: 13916–13921. doi:10.1128/JVI.01585-07

56. Zack JA, Kim SG, Vatakis DN. HIV restriction in quiescent CD4+ T cells. Retrovirology. BioMed Central; 2013;10: 37. doi:10.1186/1742-4690-10-37

57. Ospina Stella A, Turville S. All-Round Manipulation of the Actin Cytoskeleton by HIV. Viruses. 2018;10. doi:10.3390/v10020063

58. Cameron PU, Saleh S, Sallmann G, Solomon A, Wightman F, Evans VA, et al. Establishment of HIV-1 latency in resting CD4+ T cells depends on chemokine-induced changes in the actin cytoskeleton. Proc Natl Acad Sci USA. National Acad Sciences; 2010;107: 16934–16939. doi:10.1073/pnas.1002894107

59. Tardif MR, Tremblay MJ. Regulation of LFA-1 Activity through Cytoskeleton Remodeling and Signaling Components Modulates the Efficiency of HIV Type-1 Entry in Activated CD4 +T Lymphocytes. The Journal of Immunology. 2005;175: 926–935. doi:10.4049/jimmunol.175.2.926

60. Hed El A, Khaitan A, Kozhaya L, Manel N, Daskalakis D, Borkowsky W, et al. Susceptibility of human Th17 cells to human immunodeficiency virus and their perturbation during infection. J Infect Dis. 2010;201: 843–854. doi:10.1086/651021

61. Gosselin A, Monteiro P, Chomont N, Diaz-Griffero F, Said EA, Fonseca S, et al. Peripheral blood CCR4+CCR6+ and CXCR3+CCR6+CD4+ T cells are highly permissive to HIV-1 infection. J Immunol. 2010;184: 1604–1616. doi:10.4049/jimmunol.0903058

62. Prendergast A, Prado JG, Kang Y-H, Chen F, Riddell LA, Luzzi G, et al. HIV-1 infection is characterized by profound depletion of CD161+ Th17 cells and gradual decline in regulatory T cells. AIDS. 2010;24: 491–502. doi:10.1097/QAD.0b013e3283344895

63. McKinnon LR, Nyanga B, Kim CJ, Izulla P, Kwatampora J, Kimani M, et al. Early HIV-1 infection is associated with reduced frequencies of cervical Th17 cells. J Acquir Immune Defic Syndr. 2015;68: 6–12. doi:10.1097/QAI.0000000000000389

64. Stieh DJ, Matias E, Xu H, Fought AJ, Blanchard JL, Marx PA, et al. Th17 Cells Are Preferentially Infected Very Early after Vaginal Transmission of SIV in Macaques. Cell Host & Microbe. Elsevier Inc; 2016;19: 529–540. doi:10.1016/j.chom.2016.03.005

65. Cleret-Buhot A, Zhang Y, Planas D, Goulet J-P, Monteiro P, Gosselin A, et al. Identification of novel HIV-1 dependency factors in primary CCR4(+)CCR6(+)Th17 cells via a genome-wide transcriptional approach. Retrovirology. BioMed Central; 2015;12: 102. doi:10.1186/s12977-015-0226-9

66. Wacleche VS, Goulet J-P, Gosselin A, Monteiro P, Soudeyns H, Fromentin R, et al. New insights into the heterogeneity of Th17 subsets contributing to HIV-1 persistence during antiretroviral therapy. Retrovirology. BioMed Central; 2016;13: 59. doi:10.1186/s12977-016-0293-6

67. Hogg N, Patzak I, Willenbrock F. The insider’s guide to leukocyte integrin signalling and function. Nat Rev Immunol. Nature Publishing Group; 2011;11: 416–426. doi:10.1038/nri2986

68. Lee AYS, Eri R, Lyons AB, Grimm MC, Korner H. CC Chemokine Ligand 20 and Its Cognate Receptor CCR6 in Mucosal T Cell Immunology and Inflammatory Bowel Disease: Odd Couple or Axis of Evil? Front Immunol. Frontiers; 2013;4: 194. doi:10.3389/fimmu.2013.00194

69. Roederer M, Nozzi JL, Nason MC. SPICE: Exploration and analysis of post-cytometric complex multivariate datasets. Cytometry. 2011;79A: 167–174. doi:10.1002/cyto.a.21015

